# Species specific marker genes for systemic defence and stress responses to leaf wounding and flagellin stimuli in hybrid aspen and silver birch

**DOI:** 10.1101/2025.07.23.666137

**Authors:** Karlis Trevors Blums, Baiba Krivmane, Maryna Ramanenka, Roberts Matisons, Dainis Edgars Rungis, Martins Zeps, Zigmunds Orlovskis

## Abstract

In Northern Europe, climate warming is driving the northward expansion of deciduous tree species such as aspen and silver birch, while simultaneously intensifying biotic stress from pests and pathogens. This creates an urgent need for improved understanding of molecular defense mechanisms underlying stress resistance and resilience in temperate forest trees, as a basis for the development of innovative biotechnological approaches. However, progress in this area remains limited by the lack of reproducible experimental systems and well-characterized molecular markers for systemic defense responses in deciduous tree species. In this study, we aimed to identify and validate known plant defense gene markers associated with systemic stress responses in hybrid aspen and silver birch to support future functional research. Using sequence mining and phylogenetic analyses, we identified homologues of biotic stress-response genes in the genomes of both species. We then employed in vitro propagated tree clones to assess defense gene activation in distal leaves following systemic signal induction by leaf wounding and bacterial flagellin treatment at 4 and 24 hours post-induction. We identified *LOX2, MPK3*, and *EIN2* as early wounding-responsive genes in silver birch, and JAZ10 with EIN2 as potential marker genes for the combined effect of flagellin and wounding in hybrid aspen Collectively, these findings establish a reproducible *in vitro* framework for validating stress responsive genes and provide a foundation for future studies of systemic signalling, tree–microbe interactions, and resilience in ecologically and economically important forest tree species.

## 1. Introduction

Accelerating climate change, intensifying weather extremes and their legacy effects, such as outbreaks of pests and diseases (Meier *et al*., 2022; Allen *et al*., 2015), are subjecting trees to increasingly intense stresses (Seidl *et al*., 2017). These stresses can exceed the adaptive capacity (i.e., phenotypic plasticity) of local tree populations (Aitken and Bemmels, 2016), emphasizing the urgent necessity for adaptive management strategies (Sousa-Silva *et al*., 2018) to cope with growing environmental challenges (Isaac-Renton *et al*., 2018). Given the long life cycle of trees and the stresses associated with large-scale environmental changes (Moran *et al*., 2017; Seidl *et al*., 2017), effective adaptive measures should be self-sustaining and grounded in a thorough understanding of ecological, ecophysiological, and biochemical processes (Jansson *et al*., 2017). Examples of such measures include locally finetuned management (Murray *et al*., 2025; Forrester and Bauhus, 2016), as well as the inoculation of plants with beneficial microbes, such as mycorrhizal fungi, which have been suggested as promising strategies to enhance forest resilience against increasing environmental stresses (Lutter *et al*., 2023; Ciadamidaro *et al*., 2017; Bois *et al*., 2005).

In the eastern Baltic region, climate-driven shifts in forest species distribution are anticipated, with deciduous species expected to push conifers northward (Buras and Menzel, 2019). Under these conditions, the prevalence of silver birch (*Betula pendula*) and aspen (*Populus tremula*) — both currently of high commercial importance — is projected to increase (Dubois *et al*., 2020; Tullus *et al*., 2012). However, both species are sensitive to climate (Matisons *et al*., 2022; Šēnhofa *et al*., 2015), and thus, an intensification of biotic legacy effects, such as insect pest outbreaks following climatic disturbances, is also expected (Gely *et al*., 2020; Seidl *et al*., 2017). Additionally, the use of specific, highly productive genotypes — such as the most economically valuable birch clones or superior hybrid aspens — is recommended for commercial forestry (Bāders *et al*., 2024; Tullus *et al*., 2012). This underscores the need to identify innovative biotechnological approaches for improving stress tolerance in selected high yielding genotypes and to enhance understanding of tree stress responses at a molecular level (Estravis-Barcala *et al*., 2020; Bradshaw *et al*., 2019; Lelu-Walter *et al*., 2013).

Molecular stress responses have been extensively studied in herbaceous crop plants and model species such as *Arabidopsis thaliana*, however, trees with their large and complex genomes have been comparatively understudied and often limited to genus *Populus* (Harfouche, 2014). This has hitherto limited functional studies on the genetic mechanisms involved in biotic stress responses in other economically important tree species. Nevertheless, recent advances in sequencing the genomes of important forestry species such as silver birch (Salojarvi *et al*., 2017) or several aspen species (Lin *et al*., 2018) have catalysed recent progress in elucidating the molecular basis of tree responses to abiotic stresses (Estravis-Barcala *et al*. 2020). Similar to *Arabidopsis,* tree responses to chilling (frost), heat and drought, are likely mediated by abscisic acid (ABA), ROS and Ca^2+^ signalling, mitogen-activating protein kinases (MAPK), heat-shock proteins (HSP), MYB and bZIP family transcription factors (among others), resulting in antioxidant production, changes in membrane fluidity and repair (reviewed by Estravis-Barcala *et al*., 2020; Harfouche *et al*., 2014). Still, reliable stress responsive genes for studying tree responses to herbivore attack or microbial infections remain to be characterised for most tree species.

This study aimed to identify plant defence gene homologues associated with systemic stress responses in hybrid aspen and silver birch and to validate their activation using *in vitro* propagated tree clones subjected to controlled biotic stress cues. Leaf wounding and application of the immunogenic bacterial peptide flagellin were used to mimic mechanical damage caused by chewing herbivores and bacterial elicitor perception via the pattern-triggered immunity receptor FLS2 (Chinchilla, 2006), respectively. To this end, we mined the hybrid aspen and silver birch genomes for genes involved in jasmonic acid–mediated wound responses, including *LOX* and *JAZ* family members (Fürstenberg-Hägg, 2013), as well as for genes associated with salicylic acid–related defence responses to bacterial elicitors, including *FRK1, NHL10, MPK3, PAD4*, and *MYB51* (Boudsockq *et al*., 2010; Cui *et al*., 2017). Using *in vitro* propagated tree clones as a reproducible experimental platform, we characterized the activation of selected defence genes in distal leaves following induction of systemic signals by local leaf wounding and flagellin treatment at defined time points. Here, we present an *in vitro*-based framework for the initial validation of stress responsive genes of systemic defence signalling in birch and aspen, providing a foundation for future studies of intra- and inter-tree communication and biotic stress responses.

## 2. Materials and Methods

### 2.1. In-vitro propagation of tree clones

Silver birch and hybrid aspen clone cultures were obtained from the clonal collection of the LSFRI “Silava” plant physiology laboratory. For hybrid aspen *Populus tremuloides × tremula*, clone “44” was selected, as it shows superior field performance (Šēnhofa *et al*., 2016). For silver birch, clone “54-146-143”, which shows above average field performance and exceeds wild populations by 7–30 % in volume growth (Gailis *et al*., 2020; Jansson *et al*., 2017), was used. The clones were cultivated *in vitro* on 1X Murashige and Skoog (MS) media, supplemented with MS vitamins, MS micronutrients, 20 g/L sucrose, 6 g/L of Agar and 0.1 mg/L indole-3-butyric acid (IBA) at pH=5.8 (Kondratovičs *et al*. 2022).

Each clone (plantlet) was cultivated individually in a 150 mL jar filled with 15 mL of the media. During the cultivation, the jars were covered with an aluminium foil cap to prevent microbial contamination whilst enabling air flow and easy access to the plantlet for manipulation and stress treatments. To ensure controlled environmental conditions, all the clone plantlets were grown in a climate chamber maintained at 30-40% relative humidity and 25 - on four multi-layer shelf systems equipped with luminaries. All clones were grown under the same illumination with a photon flux density of 110 ± 10 µmol m^-2^ s^-1^, for the wavelengths ranging from 400 to 750 nm with 16 h light and 8 h dark photoperiod to imitate long-day conditions of growing period. For each tree species, a total of 32 plantlets were cultivated for the assessment of stress responses in orthogonal time-course experimental design, allowing four biological replicates per treatment and observation combination.

### 2.2. Stimulation of local leaves and sample collection

To assess the defence-related gene expression, the plantlets were subjected to three different treatments: a chemical stimulus with 22 amino acid fragments of bacterial flagellin (*flg22*), mechanical injury to leaves and the combination of both. These stimuli were chosen to imitate a herbivore attack (mechanical injury) or the attack of a bacterial pathogen (*flg22*) to activate defence responses related to jasmonic and salicylic acid pathways, respectively (Felix *et al*., 1999; Vlot *et al*., 2009; Fürstenberg-Hägg, 2013). The mechanical injury was applied by squeezing the leaf with tweezers in three parallel lines, while the flagellin stimulus was applied by pipetting 5 μL of 1 μM *flg22* solution (in water), on the abaxial surface of the leaf. For the combined treatment, *flg22* was applied on the wounded abaxial surface of the leaf. As the control, 5 μl of distilled nuclease free water was used.

The stimuli, including the control, were applied to the third fully unfolded leaf from the apex, marking the stimulated leaf on the outside of the jar. Plantlets were treated in a sterile environment of laminar flow before returning to the climate chamber. For the gene expression analysis, a non-stimulated (naïve) leaf - second fully unfolded leaf from the apex - distal from the treated leaf was collected 4 and 24 h after the treatment to characterise the short-term dynamics of systemic plant responses. To preserve RNA, the samples were flash-frozen in liquid nitrogen and stored at −80 °C.

### 2.3. RNA extraction and cDNA synthesis

Total RNA was extracted following a modified CTAB extraction protocol from Rubio-Piña and Zapata-Pérez (2011) adapted for use with woody plant tissue (detailed protocol in **Supplementary File 1**). The total RNA concentration and purity (260/280 and 260/230 ratios) were determined with the NanoDrop2000 spectrophotometer (Thermo Scientific). 500 ng of total RNA (260/280 and 260/230 ratios between 1.8 and 2.2) were processed for reverse transcription reaction using the Thermo Scientific Maxima First Strand cDNA synthesis kit according to manufacturer instructions. cDNA synthesis reaction was carried out using a BIO-RAD T100^TM^ thermal cycler and the reaction product was subsequently diluted to 800 μl in nuclease-free H_2_0.

### 2.4. Primer design

In order to observe potential differences in defence related gene expression in naïve systemic plant tissue (distal leaf), 8 different genes with a well described role in *Arabidopsis thaliana* defense mechanisms were chosen as the qPCR targets (**Supplementary Table 1**); *EIN2, FRK1, PAD4, LOX2, MPK3, NHL10*, *JAZ10, MYB51*, with *ACT2* chosen as a constitutively expressed reference gene for both taxa. Using gene specific protein sequences obtained from The Arabidopsis Information Resource (TAIR), BLAST was used to find potential defense gene homologues in the genomes of both tree taxa. To increase the chances of finding optimal gene homologues, BLAST search was performed in two databases for each taxon: the National Centre for Biotechnology Information (NCBI) database was used for both taxa (*Populus tremula x tremuloides taxid:47664; Betula pendula taxid:3505)*, while Comparative Genomics (CoGe) and PlantGenIE were used for birch (*Betula pendula scaffold assembly id35079, id35080*) and hybrid aspen (*Populus tremula x tremuloides v10*), respectively. The BLAST search results with the highest homology identifiers (query coverage, percent identity and e-value) were chosen as the potential homologues of each gene and subjected to reciprocal BLAST against *A. thaliana* and other taxa included in the NCBI database.

A complementary phylogenetic analysis of the selected tree homologues was performed using the MEGA (v11.0.13) Maximum likelihood algorithm (bootstrap=500). In addition to the potential tree homologues, additional *A. thaliana*, *Medicago truncatula* and *Marchantia polymorpha* sequences from NCBI and MarpolBase were included in the phylogenetic trees. Based on the *in silico* gathered data, a single potential homologue was selected for each gene as displayed in **Supplementary Figure 1** and **Supplementary Figure 2**. Using NCBI Primer-BLAST, a primer pair was designed for each selected gene sequence. All primer pairs were designed with T_m_ of around 60 □ (with ±1 □ as the maximal deviation) and PCR product size of 100-300 nt, as reported in **Supplementary Table 1**.

### 2.5. rt-qPCR analysis

The qPCR reactions were carried out using the Thermo Scientific Maxima SYBR Green/ROX qPCR Master Mix (2X) kit, with each reaction containing 2 μl of 0.3 μM primer mix, 5.5 μl of sample cDNA solution and 7.5 μl of the Maxima SYBR Green/ROX qPCR Master Mix (2X). rt-qPCR was performed on *ViiA™ 7 Real-Time PCR System (Applied Biosystems)* using 3-step cycle (initial denaturation 95°C for 10min; denaturation 95°C for 15s; annealing 60°C for 30s; extension 72°C for 30s) and analyzed with the native *QuantStudio 7 Pro 1.6.1* software. For each sample, three technical replicate qPCR reactions were carried out with each primer pair. The resulting average Ct value was then used in further calculations. Amplicons were checked for single peaks in their melting curves (**Supplementary Figure 3**) as well as visualized electrophoresis gel for the presence of single band at the expected amplicon length (**Supplementary Figure 4**).

Primer efficiency (E%) was calculated using the formula 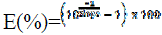, with slope being obtained by plotting the log_10_ values of 10-fold serial dilutions from 1ng to 1ag of purified cDNA (**Supplementary Figure 5**). The expression of each test gene was normalized by the expression of a single reference gene (*ACT*) using the formula ΔCt=E_ref_^Ct_ref_ / E_test_^Ct_test_, where *E_ref_* – primer efficiency of reference gene, *E_test_* – primer efficiency of test gene, taken to the power (^) of *Ct_ref_* – Ct value of reference gene, *Ct_test_* – Ct value of the test gene, respectively. In total, four expression values were obtained for each gene, for all stimuli groups and at each time-point for both birch and hybrid aspen. To determine change in defence gene expression relative to the water control, log2 fold change was calculated using the formula ΔΔCt= log2(x/y), where x - the average normalized expression (ΔCt) of a select treatment and y - the average normalized expression (ΔCt) of the control group. Analysis was done separately for each timepoint.

### 2.6. Leaf wetting experiments

Leaf wetting experiment was performed according to the method by Limm *et* al. (2009). Single leaves were cut from the plant, and petioles sealed with Parafilm to prevent water loss or entry through the petiole tip. Leaves were fully submerged into dH2O or 0,2% (w/v) rhodamineB solution or rhodamineB solution with 0,001% (v/v) Tween80. Foliar water uptake (FWU) into apoplast was calculated by weighing the mass of the detached leaf (with parafilm) and calculating using the formula FWU = (M2 − M1) − (M4 − M3), where M1-mass of leaf before submergence, M2- mass of leaf submerged for 3h then blotted on tissue to remove excess water, M3-mass of leaf after M2 measurement and air-dried for 5 min, M4-leaf after M3 measurement resubmerged for 1s, then immediately blotted on tissue to remove excess water. Leaf surface contact angle with a 5 μL drop of 1μM *flg22* was determined by taking photographs with 100mm 1X magnification macro lens at the minimal focal distance (30cm) at F2.8 and 1/60s, then analysed with the ImageJ plugin Contact Angle.jar.

### 2.7. Promoter sequence analysis

5k upstream region from TTS for AtLOX2 (AT3G45140), AtEIN2 (AT5G03280) were obtained from *TAIR* database, for BpLOX2 (Bpev01.c0523.g0011), BpEIN2 (Bpev01.c0990.g0003) form *CoGe* database and PttLOX2 (Potrx066157g26369.5), Ptt EIN2 (Potrx046169g13695.4), PpLOX2 (Potri.001G015300.1) from *Plantgenie* database. TCP4 (MA1035.1; TFmatrixID_0423), TCP20 (MA1065.1; TFmatrixID_0424), TCX5 (UN011.1) binding motifs were obtained from *JASPAR* and *PlantPan* databases. Promoter alignment and identification of transcription factor binding sites was performed with *PlantPAN 4.0* built-in *Promoter analysis tool* and *Cross species Blast2SEQ tool*.

### 2.6. Statistical analysis

To assess differences in gene expression between treatment groups, a two-way, single factor analysis of variance (ANOVA) was performed along with Tukey HSD post-hoc tests in cases of significant differences, using base R. In cases where ANOVA assumptions were violated, non-parametric Kruskal-Wallis tests and subsequent post-hoc Dunn tests were performed. Kruskal-Wallis tests were done using base R, while the “dunn.test” package (Dinno A, 2024, R package version 1.3.6.) was used for the Dunn post-hoc tests. A significance level α of 0.05 was used for all hypothesis significance tests. Heatmaps were generated using the *pheatmap* function from the “pheatmap” package (Kolde R., 2019, R package version 1.0.12) in R with centroid linkage clustering based on Euclidean distance. To assess the interaction between wound or *flg22* treatments and the different time-points of gene-expression across the 6 gene panel, a multivariate linear model was used where gene Fold Change ∼ Treatment * GeneID * Time-point. The statistical models were checked for compliance with assumptions via diagnostic plots. Data analysis was conducted in R version 4.5.0. (R Core Team). The full R-script used in statistical analysis and heatmap generation is included in the **Supplemental File 2.**

## 3. Results

### 3.1. Identification of plant stress response gene homologs

To identify plant defence gene homologs in silver birch and hybrid aspen, the BLAST hits from *Arabidopsis thaliana* query protein sequence were used in an additional reciprocal BLAST against all NCBI plant taxa. Candidate tree sequences with 99% coverage to any plant protein with a matching gene annotation were shortlisted by the employed BLAST algorithms. To confirm the homology, a phylogenetic analysis using all candidate homologues for a gene of interest (GOI) from hybrid aspen and silver birch were compared alongside two well annotated herbaceous plant (*Arabidopsis thaliana* and *Medicago truncatula)* homologues as well as liverwort *Merchantia polymorpha* homologue as an outgroup (**Figure 1A**). For each GOI, hybrid aspen and silver birch sequences with highest coverage and identity to *A. thaliana* and *M. truncatula* and different from *M. polymorpha* were selected for primer design (**Supplementary Table 1**). Primer pairs displaying a single-peak melting curve, as well as yielding the desired amplicon (**Supplementary Figure 3A,B**) were further tested for their PCR efficiency (E%) which typically ranged between 90-110% (**Supplementary Table 1**). *ACT2* homologues in silver birch and hybrid aspen were selected as a reference genes as they typically displayed earlier amplification (Ct=18) compared to *TUB*, *SAND* or *GAPDH* (**Supplementary Figure 3C**) as well as did not display any significant differences in amplification cycle across water, *flg22,* wound and wound+flg22 treatments in silver birch at 4h (F=1,245; p=0,337) and 24h (F=0,64912; p=0,6101) or hybrid aspen at 4h (F=1,247; p=0,336) and 24h (F=1,9681; p=0,2345). To confirm whether normalized expression results obtained with a single (*ACT2*) reference gene would differ from using multiple references, we performed a control check with random test genes and treatments using one additional reference gene candidate for each tree species (**Supplementary Figure 5**). Since similar results were obtained with either a single or two reference gene normalization, the downstream analysis for all samples was performed with a single *ACT2* reference at 4h and 24h after stimulation of a local leaf. Time-point and species specific expression trends were observed such as induction of EIN2 in distal leaves upon local wounding in Silver birch or suppression in hybrid aspen.

**Figure 1.**
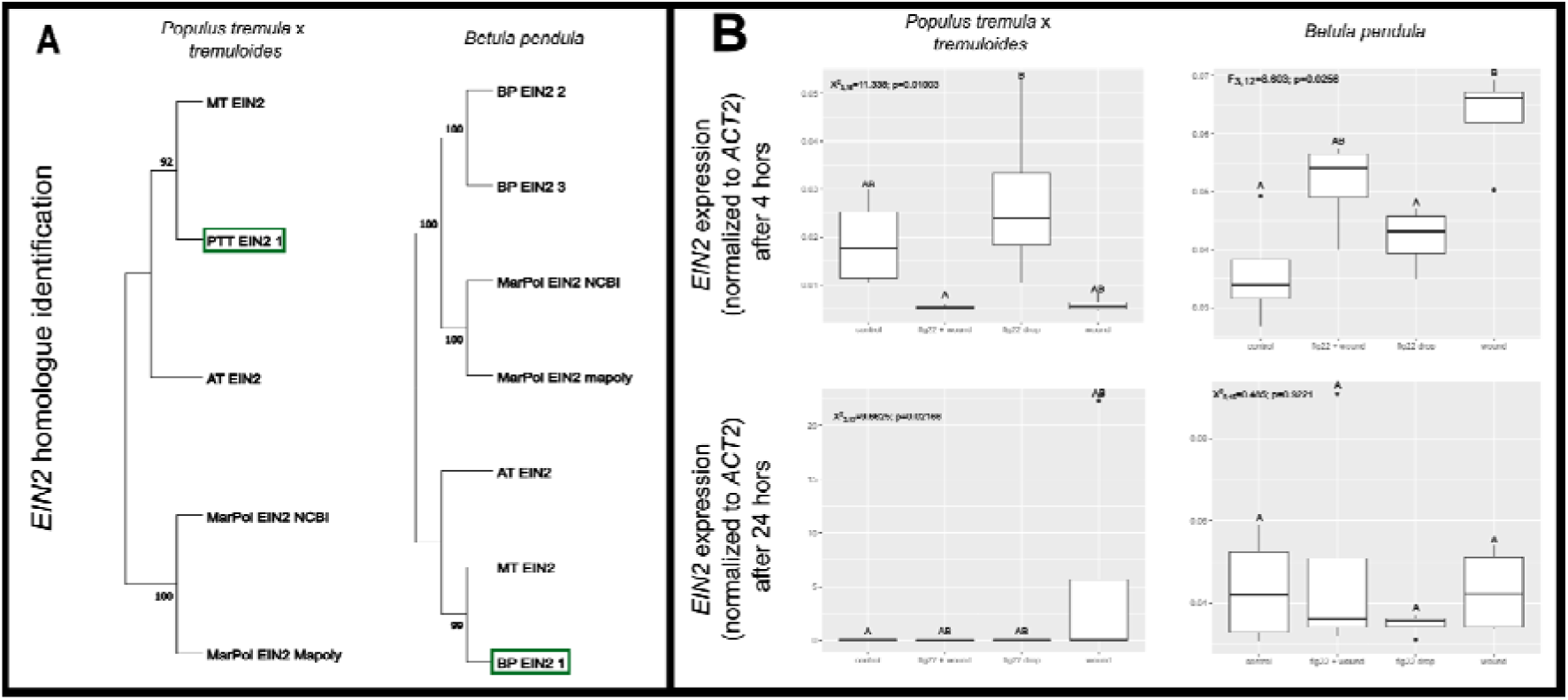
Example identification of *EIN2 (ETHYLENE INSENSITIVE 2)* homologue in silver birch and hybrid aspen for downstream characterisation of systemic intra-plant biotic stress responses. (**A**) Phylogenetic analysis of *EIN2* homolog from *Betula pendula* (BP) (left) and *Populus tremula* x *tremuloides (PTT)* (right). Green rectangles indicate the sequences selected for primer design. The trees are based on Maximum likelihood (bootstrap) and include the homologous sequences from *A. thaliana (AT)*, *M. truncatula* (MT) and *M. polymorpha* (MarPol). Phylogenetic trees for all other genes are displayed in **Supplementary Figure 1** (BP) and **Supplementary Figure 2** (PTT). (**B)** Relative expression of *EIN2* four and 24h post stimulation with flg22, wounding or their combination. Ct values were calculated as average from 3 technical replicate measurements for each sample. Gene expression was normalized to ACT2 as reference (ΔCt). Three test genes and ACT2 reference were measured together in the same qPCR run for 4 independent biological replicates of at least 2 treatments. The values for all gene ΔCt are available in **Supplementary Table 2**. Letters above boxplots indicate statistically significant differences between treatments, determined with two-tailed ANOVA or Kruskal-Wallis tests, with subsequent post-hoc tests.

### 3.2. Characterisation of stress responsive gene expression in systemic leaves

The tested wounding and *flg22* stimuli and their combination demonstrated significantly different effects (relative to the control) on gene expression patterns in silver birch (p<0.01), as well as hybrid aspen (p=0.03), indicating clear inducible systemic responses in the distal leaves (**Supplementary Table 3**). The gene expression patterns showed dependency on time after stimulation of the local leaf, highlighting species specific early and late gene response to biotic stress (**Figure 2**). Birch showed significant early induction of *LOX2* (log_2_ fold change = 2.98; p=0.03), *MPK3* (log_2_ fold change = 1.73; p<0.01) and *EIN2* (log_2_ fold change = 0.81; p=0.03) 4h after wounding. The effect, however, was clearly transitory as it disappeared after 24h. In contrast, *flg22* treatment did not display any significant gene induction in systemic tissues neither 4 h nor 24 h after stimulation of the local leaves. Four hours after the application of the combined wounding and *flg22* treatment only *MPK3* expression was significantly elevated (log_2_ fold change =1.34; p=0.03), indicating potential antagonistic responses to wounding and *flg22* treatment in silver birch.

**Figure 2.**
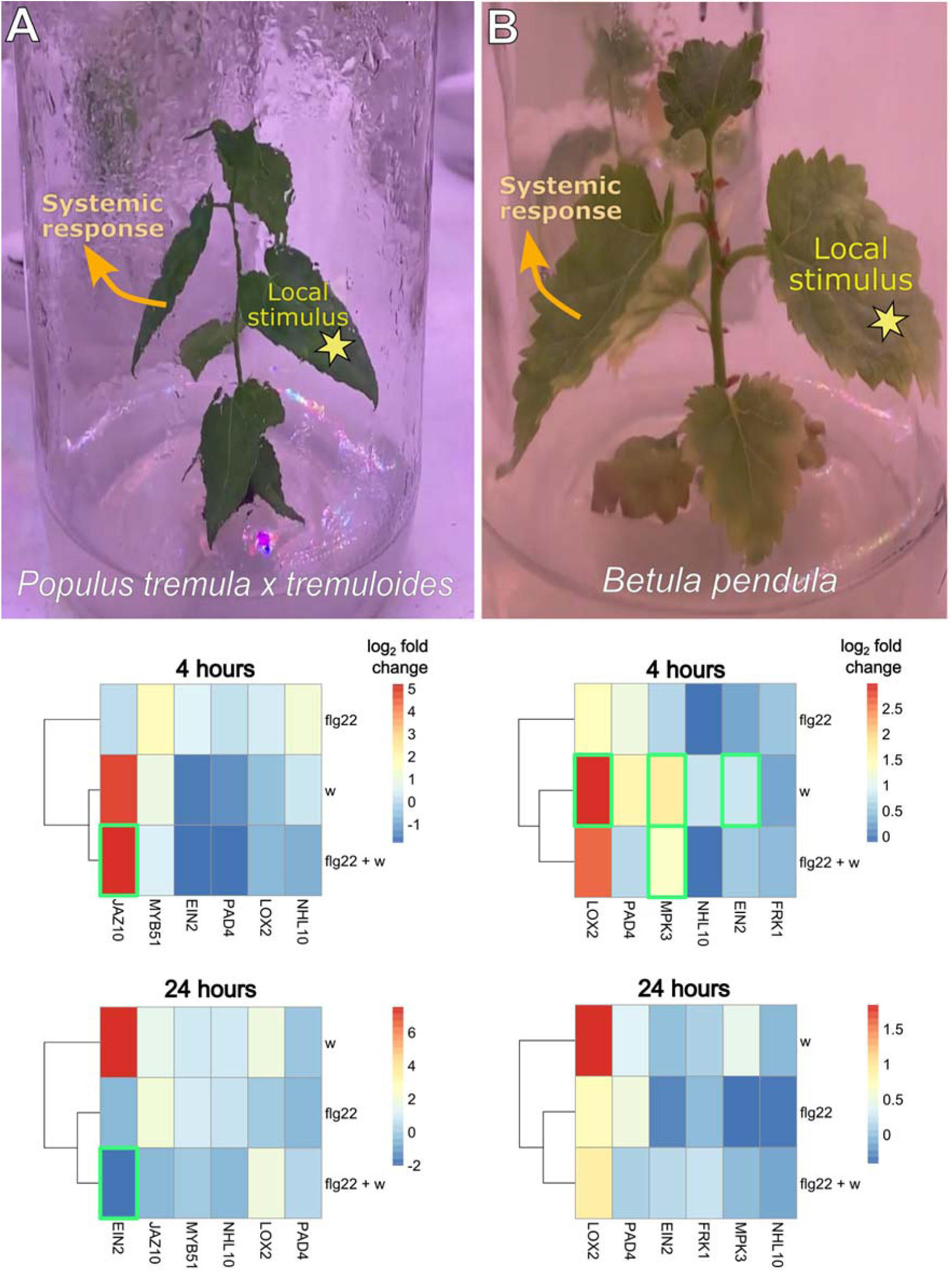
Experimental setup and relative gene expression in systemic leaves of hybrid aspen (column A) and silver birch (column B) clones in response to wounding and *flg22* treatment. The stress stimuli were applied to a local leaf and systemic responses were measured in a distal leaf of *in vitro* grown tree clones. The heatmaps demonstrate log2 fold-changes in gene expression (x-axis) for each treatment group (y-axis) relative to the water control at 4 hour and 24 hours post stimulation of a distal leaf. The green rectangles indicate statistically significant (p<0.05) differences in gene expression in systemic leaves based on ANOVA post-hoc tests. Local leaf stimuli are *flg22* - 5 µL of 1µM flagellin; *w* - forceps wounding; *flg22 + w* - combination of the flg22 with wounding. Response similarity to treatments is based on centroid linkage and clustering by Euclidean distance. The fold change is calculated using formula ΔΔCt=log2(x/y), where x - the average normalized expression (ΔCt) of a select treatment and y - the average normalized expression (ΔCt) of the control group. ΔCt is calculated as gene expression normalized to *ACT2* as reference. Full normalized expression data for each gene along with baseline expression in water control treatments are displayed in **Supplementary Figure 7.** Source data for calculating the fold-change values are available in **Supplementary Table 2.**

In contrast to silver birch, hybrid aspen showed significant interaction between treatment type, identity of the activated genes and timepoints after stimulation (p<0.001) (**Supplementary Table 3**), indicating complex control. Only the combined wounding and *flg22* treatment induced significant aspen gene activation at 4 h and 24 h (**Figure 2**). *JAZ10* was significantly (log_2_ fold change =5.18; p=0.01) induced 4h after the combined wounding and *flg22* treatment and displayed increased expression relative to the water control in the wounded plants as well, albeit not significantly. Surprisingly, the known wound-responsive JA pathway marker gene *LOX2* showed no significant induction in hybrid aspen at neither 4 h nor 24 h after the stimulation. The ethylene signalling gene *EIN2* displayed significant suppression (log_2_ fold change =-2.03; p=0.01) 24 h after the combined wounding and *flg22* treatment. Still, it demonstrated a tendency to be induced after wounding, suggesting potential opposing effects of *flg22* and wounding in the systemic responses of hybrid aspen leaves (**Figure 2**).

### 3.3. Promoter divergence in stress responsive genes of birch and aspen

Given the species-specific responses of the measured genes to wounding and *flg22* treatment, we hypothesised that promoter divergence may in part explain the different regulation of genes such as *LOX2* and *EIN2* in response to the same wounding or *flg22* stimulus. Alternatively, we also considered the possibility that droplet treatments containing *flg22* may have differentially penetrated the leaf surface of silver birch and hybrid aspen and perhaps contributed to the different responses to *flg22*. To explore these possibilities, we first performed additional experiments to determine droplet contact angles (Aryal & Neuner, 2010) with the leaf surface as well as measured leaf wetting by submergence in water (Limm *et al.,* 2008) as well as rhodamine B dye to enable contrast microscopy. Interestingly, we found that leaves of silver birch demonstrate average contact angles of 5 µL flg22 droplet to be <90° and in hybrid aspen <30°, indicative of good wetting properties, according to Aryal & Neuner (2010) (**Supplemental Figure 8A**). Next, we submerged the leaves for 3 h to determine water uptake from leaf surface into the apoplast and concluded that both species display comparable wetting characteristics (**Supplemental Figure 8B**). Furthermore, the application of 5 µL rhodamine B droplet confirmed penetration of dye into mesophyll of both aspen and birch (**Supplemental Figure 8C**). While the saturation was relatively higher and more uniform across aspen leaves compared to patchy dye penetration across birch leaves, the stomata openings appear to serve as passage for rhodamine B into the mesophyll of birch leaves (**Supplemental Figure 8C,** row 4). Given that *flg22* can show activity in nM concentration range (Felix *et al*., 2002) and we applied *flg22* at 1µM concentration, it may be unlikely that the observed differences in gene expression between species could be attributed to differential dosage of *flg22* in the apoplast or lack of droplet uptake by the leaf surface.

To characterise the potential evolutionary divergence of tree promoter sequences that may contribute to differential binding of transcription factors to the measured gene regulatory sequences, we performed additional *in silico* analysis of silver birch and hybrid aspen *LOX2* putative promoter sequence and TCP transcription factor (TF) binding sites within 5k region upstream UTR and TSS (**Figure 3A**). TCP TFs are known regulators of JA synthesis in plant immunity and development (Lopez *et al*., 2015). *Populus tremula x tremuloides* and *Populus trichocarpa* demonstrated good synteny of conserved regions throughout the *LOX2* promoter (**Figure 3B**). In contrast, hybrid aspen and silver birch showed comparatively short conserved fragments (12-20nt) rearranged throughout the 5k promoter region (**Figure 3C**). Moreover, the conserved promoter regions (e.g., region 9) displayed differences in predicted TF binding sites (**Figure 3C**). To further verify this, we selected binding motifs of known *LOX2* transcriptional regulators - positive regulator *TCP4*, negative regulator *TCP20,* and their interactor TCX8 (**Figure 3D**). Since the TF binding matrices of TCP and TCX factors are only available for *Arabidopsis thaliana* homologs, we used this only as indicative proxy analysis for their putative correspondence to *LOX2* and *EIN2* from hybrid aspen and silver birch. The two genes were selected because they originally showed tree-specific differences in expression upon wounding or WF stimuli (**Figure 2**). As control, we also included *AtLOX2* and *AtEIN2* promoter sequences. Interestingly, TCP4 binding sites were found in both *BpLOX2* and *BpEIN2* but not in the respective hybrid aspen homologs (**Figure 3E**). TCP20 binding sites were found only in *PttLOX2* but not *BpLOX2* promoter (**Figure 3F**). Interestingly, the TCP4 and TCP20 binding motifs were found in EIN2 promoter regions in either of the two tree species but not *A. thaliana*, potentially suggesting that downstream gene responses to TCP activating signals (such as wounding) may be different in *Arabidopsis* and hybrid aspen or silver birch. TCX8 - a known interactor of class I and II TCPs and DREAM complex (Noh *et al.,* 2021) - displayed numerous but distinct binding sites along the promoter regions of *LOX2* and *EIN2* from all three species (**Figure 3G**), further highlighting putative complex regulatory circuits that could underlie the species specific responses to wounding and *flg22* based on the observed promoter divergence (**Figure 3C**).

**Figure 3.**
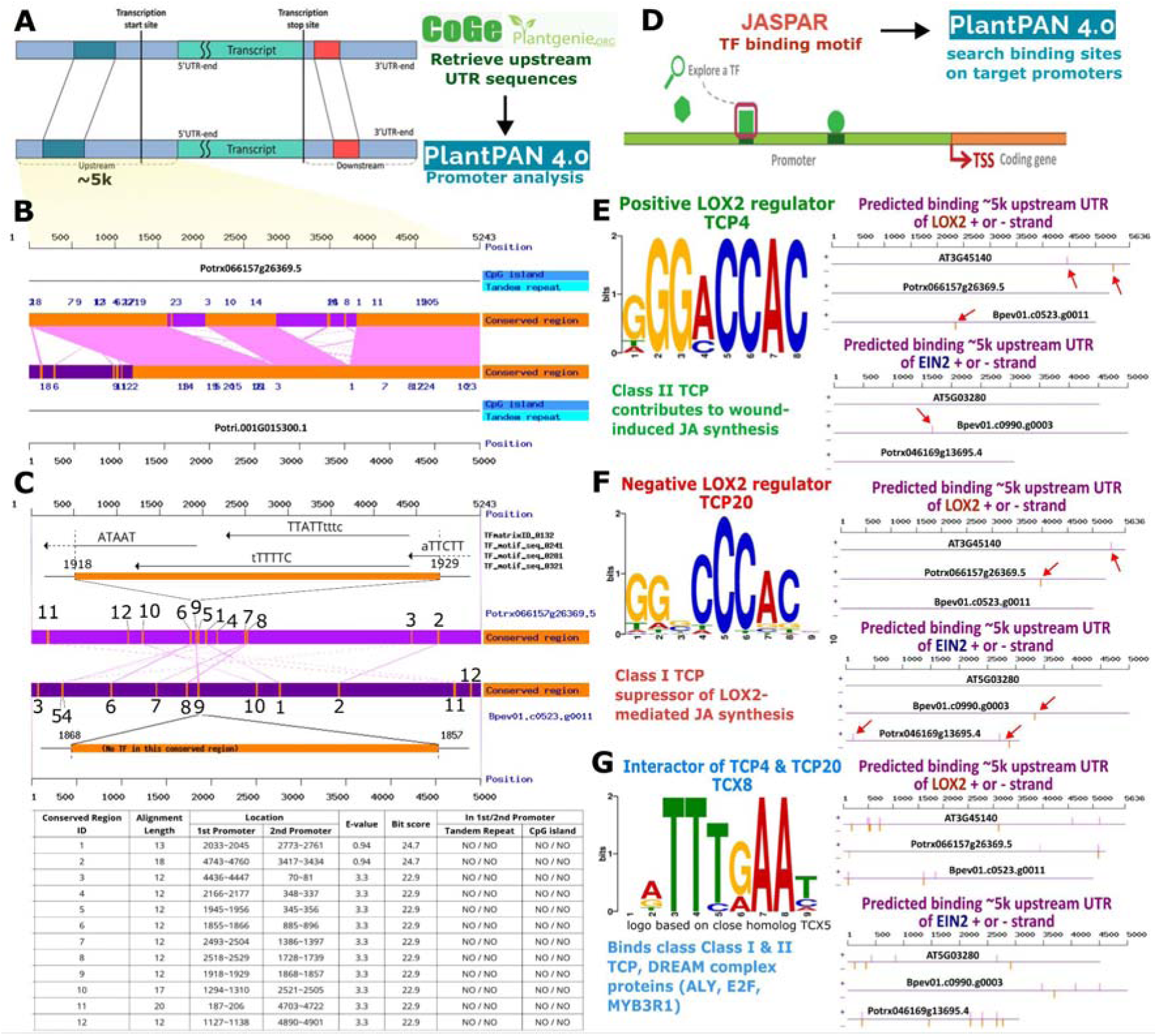
Silver birch and hybrid aspen display sequence divergence in *LOX2* promoter region and putative transcription factor binding sites. 5k upstream UTR sequences of *LOX2* gene were retrieved from the *CoGe* and *Plantgenie* databases and aligned with *PlantPan v4.0* to identify conserved regions and transcription factor (TF) binding motifs in the putative promoter region of *LOX2* gene (**A**). Promoter regions of *LOX2* homolog from *Populus tremula x tremuloides* (Ptt, Potrx066157g26369.5) displays higher sequence conservation and synteny with *Populus trichocarpa* (Pt, Potri.001G015300.1) (**B**) compared to more divergent promoter and putative TF binding sites in *Betula pendula* (Bp, Bpev01.c0523.g0011) (**C**). Only 12 conserved regions of 12-20 nt in length were identified in the 5k upstream UTR region of *Bp* and *Ptt* corresponding to different locations within the putative promoter region. As an example, region 9 from *Ptt* and *Bp* displays different TF binding sites from the available *PlantPan* annotations (**C**). Binding matrices of known *LOX2* regulators - TCP and TCX transcription factors - were acquired from JASPR database for their specific correspondence to *LOX2* and *EIN2* promoter regions from *Arabidopsis thaliana* (*At*), *Ptt* and *Bp* using *PlantPan v4.0 promoter analysis tool* (**D**). TCP4, TCP20 and TCX8 binding sites are displayed on the + and - strand of *LOX2* and *EIN2* promoter regions (red arrows) and compared across the three plant species (**E-G**). Only 3k upstream UTR region of *EIN2* homolog from *Ptt* (*Potrx046169g13695.4*) was retrieved due to close proximity of transcription stop site of the upstream gene.

## 4. Discussion

### 4.1. In vitro tree clones as models for plant systemic signalling research

The present study is the first to our knowledge to employ *in vitro* propagated tree clones as a model for studying intra-tree responses to stress signals and offers a scalable screening platform for design and discovery of tree stress response genes to diverse biotic stressors and their combinations. The applied methods provide certain potential advantages, including rapid propagation time, genetic uniformity due to clonal propagation, controlled growth and media conditions, sterile environment absent from uncontrolled pest or pathogen infections and the ability to precisely subject trees to selected defence elicitors or individual stress elements, potentially overcoming many challenges in future elucidation of molecular mechanisms for tree biotic and abiotic stress tolerance associated with heterogeneity and interaction of multiple environmental and stress factors in greenhouse and field studies (Estravis-Barcala, 2020; Harfouche *et al*., 2014). Early identification of stress-responsive genes using *in vitro* systems would accelerate follow-up functional studies in soil-grown saplings under growth chamber, greenhouse, or field conditions, enabling a wide range of biological investigations.

The experimental system herein allowed for optimal selection of tree stress inducing treatments and their response time estimation. For example, we discovered that only the combined treatment of wounding and *flg22* application to local leaves triggered significant systemic activation of JA-response gene *JAZ10* after 4h in hybrid aspen (**Figure 2**), while the individual wound and *flg22* treatments failed to induce consistent early responses. In contrast, leaf wounding displayed more early responding genes compared to the other treatments in silver birch. Selection of reliable and reproducible stress inducing treatments and their marker genes could be a key primer for several applications such as mechanistic studies on inter-tree signals and responses (Orlovskis *et al*., 2024) in the future.

### 4.2. Species specificity of SA or JA responses upon wounding or flg22 application

While certain marker genes of early responses in pattern triggered immunity (PTI) have been well described in *Arabidopsis* such as early response genes *NHL10 (NDR1/HIN1-LIKE 10*) or MAPK-specific target gene *FRK1 (FLG22-INDUCED RECEPTOR KINASE1)* (Boudsocq *et al*., 2010), no significant gene expression differences compared to the water control treatment were observed for these genes in *flg22* treatment in silver birch or hybrid aspen, potentially indicating different timing or species specific differences in the mechanisms of early FLS2-mediated PTI signalling and their responses. These can be attributed to species-specific differences in the recognition of shortened *flg22* ligands by FLS2 receptor-co-receptor complexes (Mueller *et al.,* 2012) or, alternatively, heterogeneity of flagellin fragments (Colaianni *et al.,* 2021) produced by leaf endophyte communities that may interfere with responses to exogenously applied pathogenic flagellin.

Alternative explanation for species specific effects would be differential penetration of *flg22* through the cuticle and leaf surface of hybrid aspen and silver birch. While hybrid aspen leaves displayed greater wettability than silver birch, the water and dye solution uptake by leaves during the 3h period was comparable in both species and not enhanced by addition of surfactant (**Supplemental Figure 8**), corroborating the capability of foliar solute uptake in trees (Schreel & Steppe, 2020). Furthermore, application of 5 µL dye droplet comparable to the *flg22* treatment suggests more uniform uptake across the leaf surface of aspen but more patchy entry into the apoplast via open stomata in birch. While we did not determine the penetration of water or *flg22* via the wound sites in comparison to passive diffusion across leaf surface, inclusion of water + wound control could also test for the possibility of dilution of DAMPs (e.g., cAMP, cellobiose) by the added 5µL solution. Thus, future experiments may better distinguish whether interplay between wound and *flg22*-induced signals may be associated with downstream hormonal crosstalk or differences in elicitor concentration and entry into the apoplast.

*PAD4 (PHYTOALEXIN DEFICIENT4)* and *EDS1 (ENHANCED DISEASE SUSCEPTIBILITY1)* constitute a well-established module in PTI (Pruitt *et al*., 2021), as well as abiotic stress (Szechynska-Hebda *et al*., 2016) signalling and a positive regulator of salicylic acid (SA) pathway in *Arabidopsis* (Wiermer *et al*., 2005). However, *PAD4* did not display significant response to *flg22* or combined wound and *flg22* treatments of the tree species, suggesting a potentially different *flg22* perception and signalling mechanisms than *Arabidopsis*. Both *PAD4* and plant transcription factor *MYB51* are also involved in glucosinolate and callose metabolism in innate immune responses to *flg22* (Bednarek *et al*., 2009; Clay *et al*., 2009) and mediate plant responses to phloem feeding insects (Kim & Jander, 2007; Luis & Shah, 2015). When tested in hybrid aspen, *MYB51* did not show any significant responses to wounding or *flg22* treatments.

*LOX2 (LIPOXIGENASE2)* encodes a key enzyme in jasmonic acid (JA) biosynthesis (Wasternack *et al*., 2013) while *JAZ10 (JASMONATE ZIM-Domain10)* is an important component in downstream JA signalling (Chini *et al*., 2007) in wound responses and control of glucosinolate production in *Arabidopsis* (Fürstenberg-Hägg *et al*., 2013). Interestingly, *LOX2* displayed induction in silver birch but not in hybrid aspen (**Figure 1**), indicating species specific responses or response timing to wounding. However, *JAZ10* displayed upregulation upon wounding as well as the combined wounding and *flg22* treatment in aspen. Since the JA and SA pathways are considered antagonistic (Gimenez-Ibanez & Solano, 2013), this may potentially illustrate a more complex interaction between SA and JA pathways in hybrid aspen compared to *Arabidopsis*.

*MAPK (MITOGEN ACTIVATED PROTEIN KINASES)* are important regulators of herbivory associated local wounding as well as plant systemic responses (Hettenhausen *et al*., 2015). MPK3 was induced in systemic tissues upon distal wounding and *flg22* treatment in silver birch (**Figure 2**). *MPK3* and *MPK6* are known activators of ethylene (ET) biosynthesis in *Arabidopsis* (Liu & Zhang, 2006; Broekgaarden *et al*. 2015). As reviewed in Broekgaarden *et al*. (2015), ET signalling also regulates *flg22*–triggered PTI and hormonal crosstalk between SA-mediated responses to biotrophic pathogens and JA-mediated responses to wounding and chewing herbivores. *ETHYLENE INSENSITIVE2 (EIN2)* is an important ET signalling component (Yoo *et al*., 2009) and was induced in silver birch upon wounding, consistent with the observed *LOX2* and *MPK3* responses and involvement in wound signalling (Broekgaarden *et al*. 2015).

Interestingly, the opposite regulation of *EIN2* during wounding alone versus combined wounding and *flg22* treatment in hybrid aspen may indicate crosstalk between the *flg22*-dependent SA pathway and the wound-inducible JA pathway. Moreover, the culture medium used in our experiments contained 0.5 µM IAA (auxin), which is required for rooting and propagation of clonal cuttings but is also known to interact with other phytohormone pathways, including suppression of SA-mediated host defences and promotion of bacterial virulence (Djami-Tchatchou et al., 2020; McClerkin et al., 2018; Mutka et al., 2013). Consequently, systemic stress responses observed under in vitro conditions may differ in soil-grown plants and warrant further investigation under more natural growth substrates.

Finally, as this study focused on a single commercially grown, high-yielding clone per species, future work should assess stress-responsive gene expression across a broader range of genotypes. Early identification of such markers using in vitro systems would facilitate targeted functional validation in soil-grown saplings under growth chamber, greenhouse, and field conditions, thereby supporting diverse downstream biological investigations.

## Conclusion

We have identified *BpLOX2, BpMPK3*, and *BpEIN2* as suitable early response marker genes for wounding in silver birch, while *PttJAZ10* and *PttEIN2* responded to a combination of wounding and flagellin treatment in hybrid aspen. The selection of biotic stress treatments and the identification of response marker genes will contribute to future research on systemic stress responses both within and between trees, using *in vitro* and soil-propagated clones. This system offers a valuable platform for exploring the role of different microbial inoculants and the cross-communication between tree species or intra-specific genotypes. Specifically, it will facilitate investigations into the impact of pathogen attacks on trees interconnected by common mycorrhizal networks (CMN), helping to uncover the mechanisms behind CMN-mediated defense responses in forest ecosystems.

## Author Contributions

Conceptualization: ZO; methodology: KB, BK, MR, ZO; investigation: KB, ZO, MB, BK; formal analysis: KB, ZO, RM; data curation: KB, ZO; writing - original draft preparation: ZO, KB; writing - review and editing: ZO, KB, RM; visualization: ZO, KB; supervision: ZO, DR, MZ; project administration: ZO, MZ; funding acquisition: ZO. All authors have read and agreed to the published version of the manuscript.

## Conflicts of Interest

The authors declare no conflicts of interest.

## Supplemental Tables

**Table S1.**
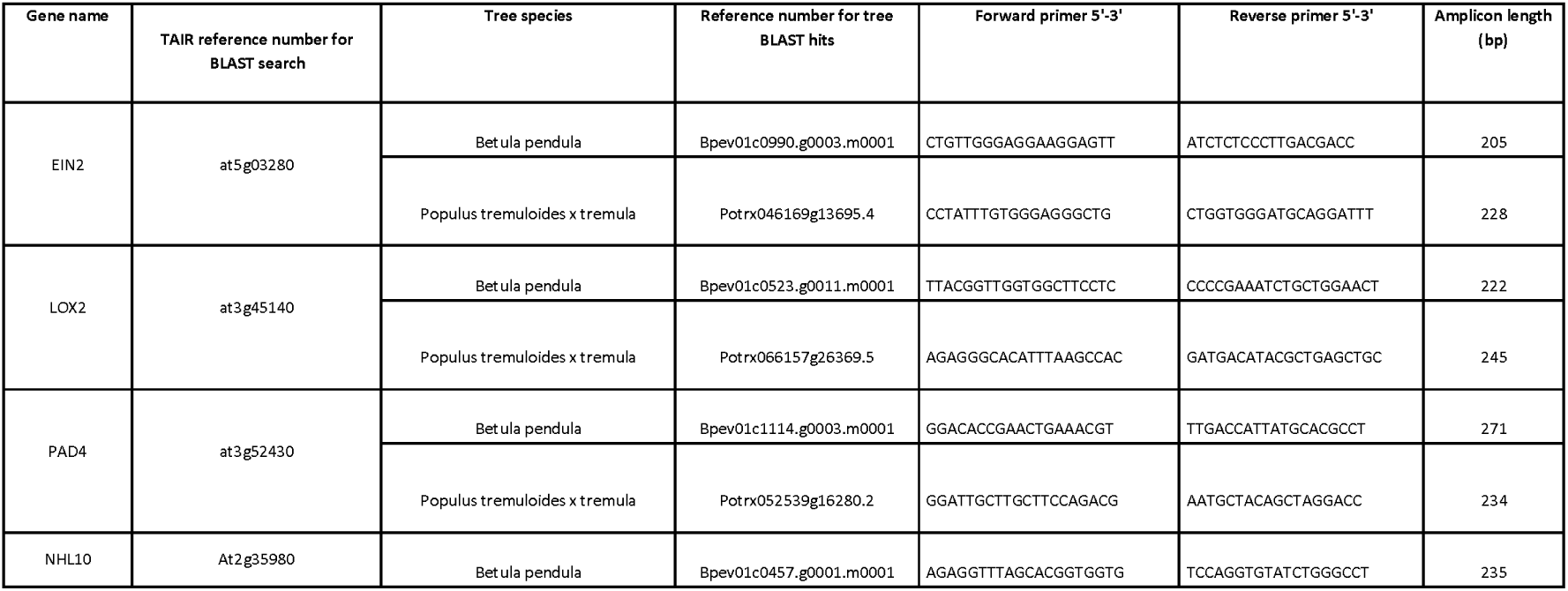
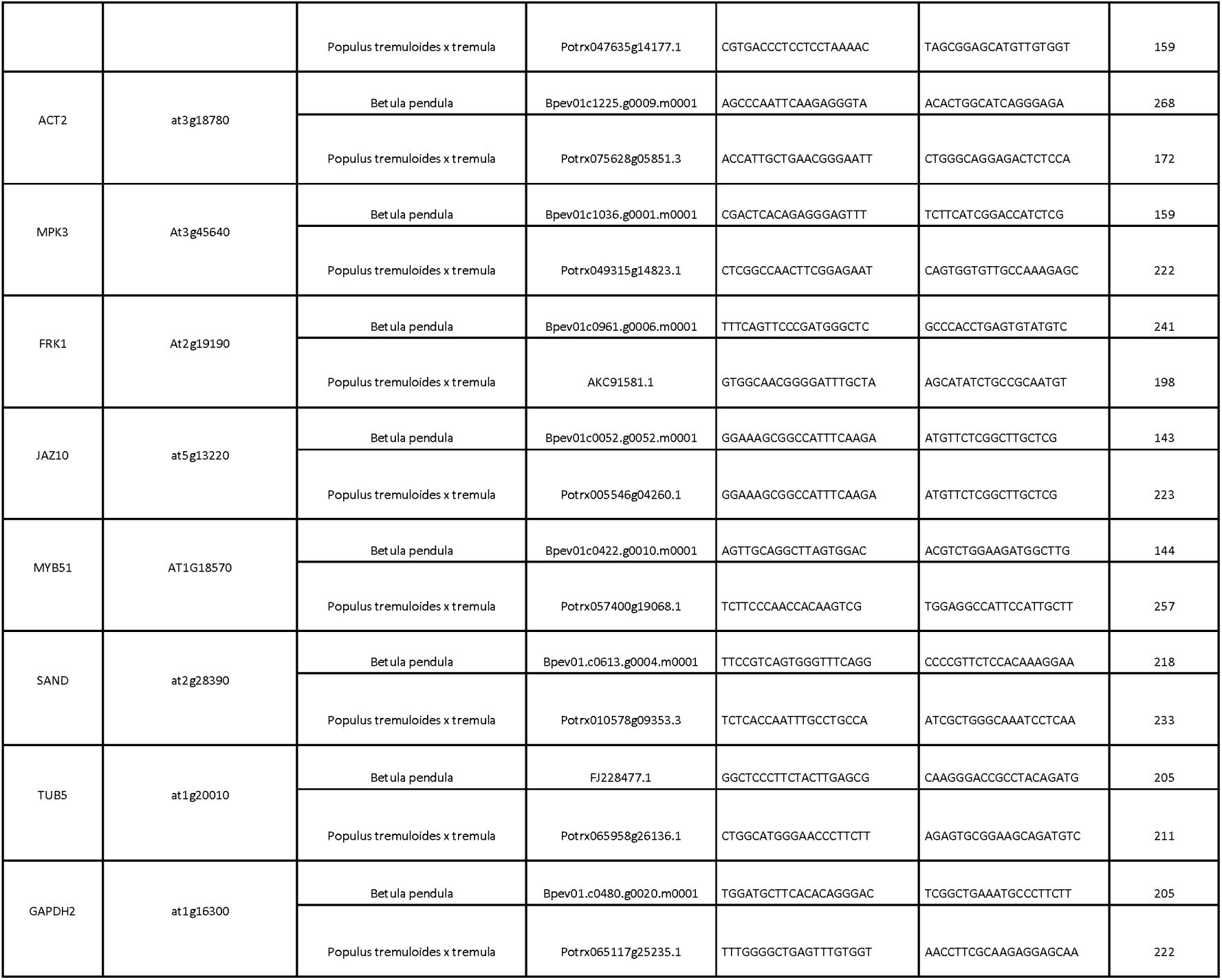
List of silver birch and hybrid aspen stress response genes used in the study. Primer sequences are provided alongside the corresponding gene ID from TAIR, CGe, PlantGene databases. Amplification factor was calculated based on 10-fold template DNA dilution series ranging from 100ag to 1ng and displayed for all genes used in **Figure 1**. Genes that did not display a specific single band in the electrophoresis and did not produce consistent melting curves (**Supplementary Figure 3**), the amplification factor was not calculated.

**Table S2. Source data table (.xls) with ΔCt and ΔΔCt values used to generate expression graphs in Figure 1B, heatmaps in Figure 2 and expression boxplots in Supplemental Figure 5.** The expression of each test gene was normalized by the expression of reference gene (ACT) using the formula ΔCt=Eref^Ctref / Etest^Cttest, where Eref – primer efficiency of reference gene, Etest – primer efficiency of test gene, taken to the power (^) of Ctref – Ct value of reference gene, Cttest – Ct value of the test gene, respectively. log2 fold change was calculated using the formula ΔΔCt= log2(x/y), where x - the average normalized expression (ΔCt) of a select treatment and y - the average normalized expression (ΔCt) of the control group.

**Table S3.** Generalized linear model results for testing the differences in gene expression between treatments across the panel of genes used in Figure 1. The model tests for the interaction of wound or *flg22* treatments with 4h and 24h timepoints and individual expression patterns of each gene (Fold Change ∼ Treatment * GeneID * Time-point).

| Silver birch | Df | Sum Sq | Mean Sq | F value | Pr(>F) | Significance |
| --- | --- | --- | --- | --- | --- | --- |
| Treatment1 | 2 | 8,428 | 4,2139 | 5,4797 | 0,00545 | ** |
| Gene | 5 | 9,33 | 1,866 | 2,4266 | 0,04002 | * |
| Timepoint | 1 | 26,904 | 26,9036 | 34,9851 | 4,20E-08 | *** |
| Treatment1:Gene | 10 | 6,048 | 0,6048 | 0,7865 | 0,64162 |  |
| Treatment1:Timepoint | 2 | 0,677 | 0,3385 | 0,4401 | 0,64513 |  |
| Gene:Timepoint | 5 | 9,334 | 1,8668 | 2,4275 | 0,03995 | * |
| Treatment1:Gene:Timepoint | 10 | 2,668 | 0,2668 | 0,347 | 0,96565 |  |
| Residuals | 105 | 80,745 | 0,769 |  |  |  |
| Adj. R2 | 0,2531 |  |  |  |  |  |
| Hybrid aspen | Df | Sum Sq | Mean Sq | F value | Pr(>F) | Significance |
| Treatment1 | 2 | 15,368 | 7,6839 | 3,7558 | 0,02661 | * |
| Gene | 5 | 78,781 | 15,7561 | 7,7013 | 3,41E-06 | *** |
| Timepoint | 1 | 0,037 | 0,0367 | 0,0179 | 0,89371 |  |
| Treatment1:Gene | 10 | 32,845 | 3,2845 | 1,6054 | 0,11515 |  |
| Treatment1:Timepoint | 2 | 6,501 | 3,2503 | 1,5887 | 0,20911 |  |
| Gene:Timepoint | 5 | 68,361 | 13,6722 | 6,6827 | 1,95E-05 | *** |
| Treatment1:Gene:Timepoint | 10 | 112,282 | 11,2282 | 5,4881 | 1,63E-06 | *** |
| Residuals | 104 | 212,774 | 2,0459 |  |  |  |
| Adj. R2 | 0,4603 |  |  |  |  |  |

## Supplemental Figures

**Supplementary Figure 1.**
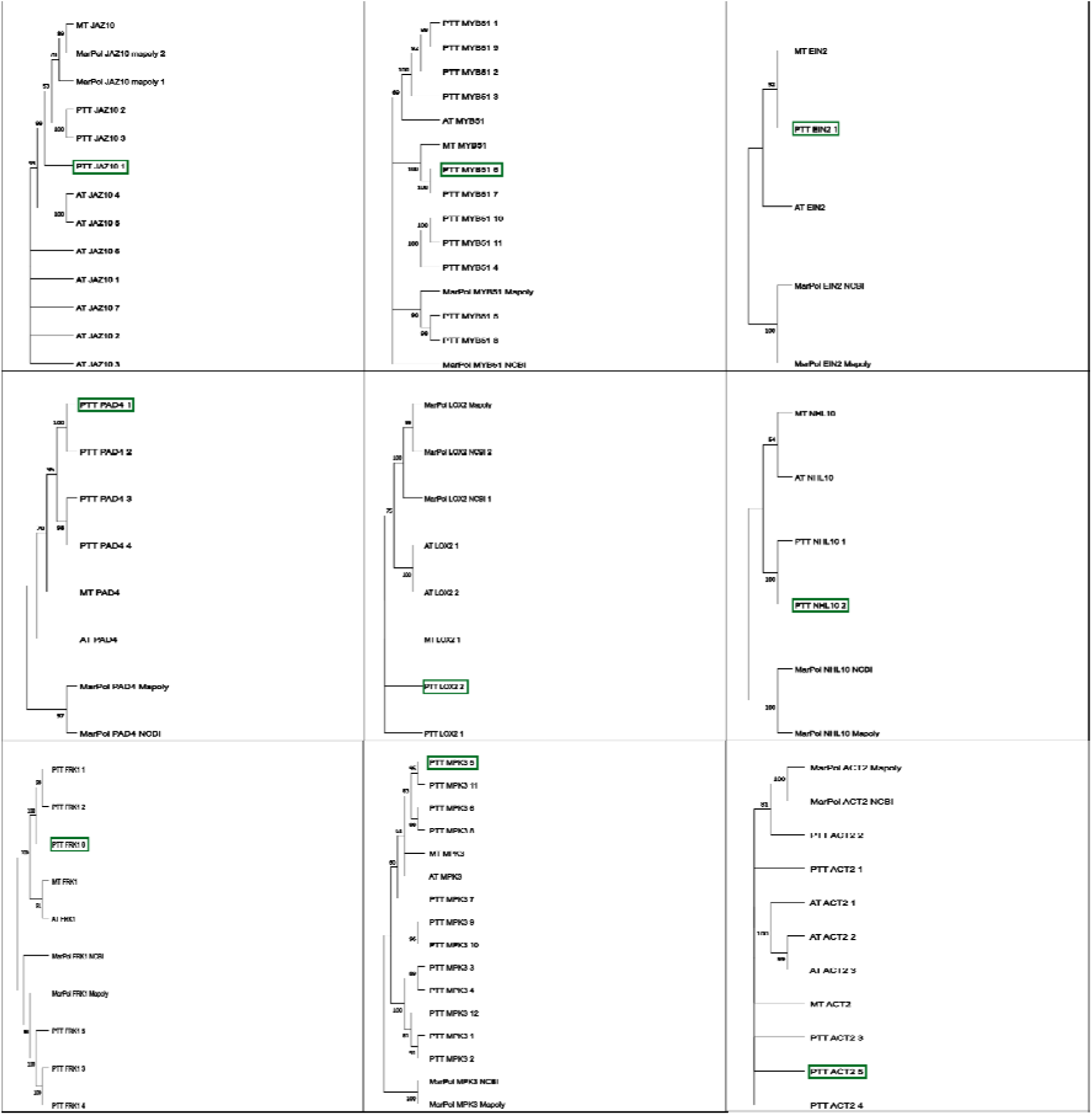
Cladograms of putative *Populus tremula x tremuloides* marker gene homologs. Phylogeny was based on translated protein sequences of *JAZ10, MYB51, EIN2, PAD4, LOX2, NHL10, FRK1, MPK3*, and reference gene *ACT2* from *Populus tremula x tremuloides* (PTT), *Arabidopsis thaliana* (AT), *Medicago truncatula* (MT), *Marchantia polymorpha* (MarPol). In case of multiple spliced variants, all sequences were included for tree construction using the Maximum likelihood method. Green box denotes the homolog used for primer design in the study.

**Supplementary Figure 2.**
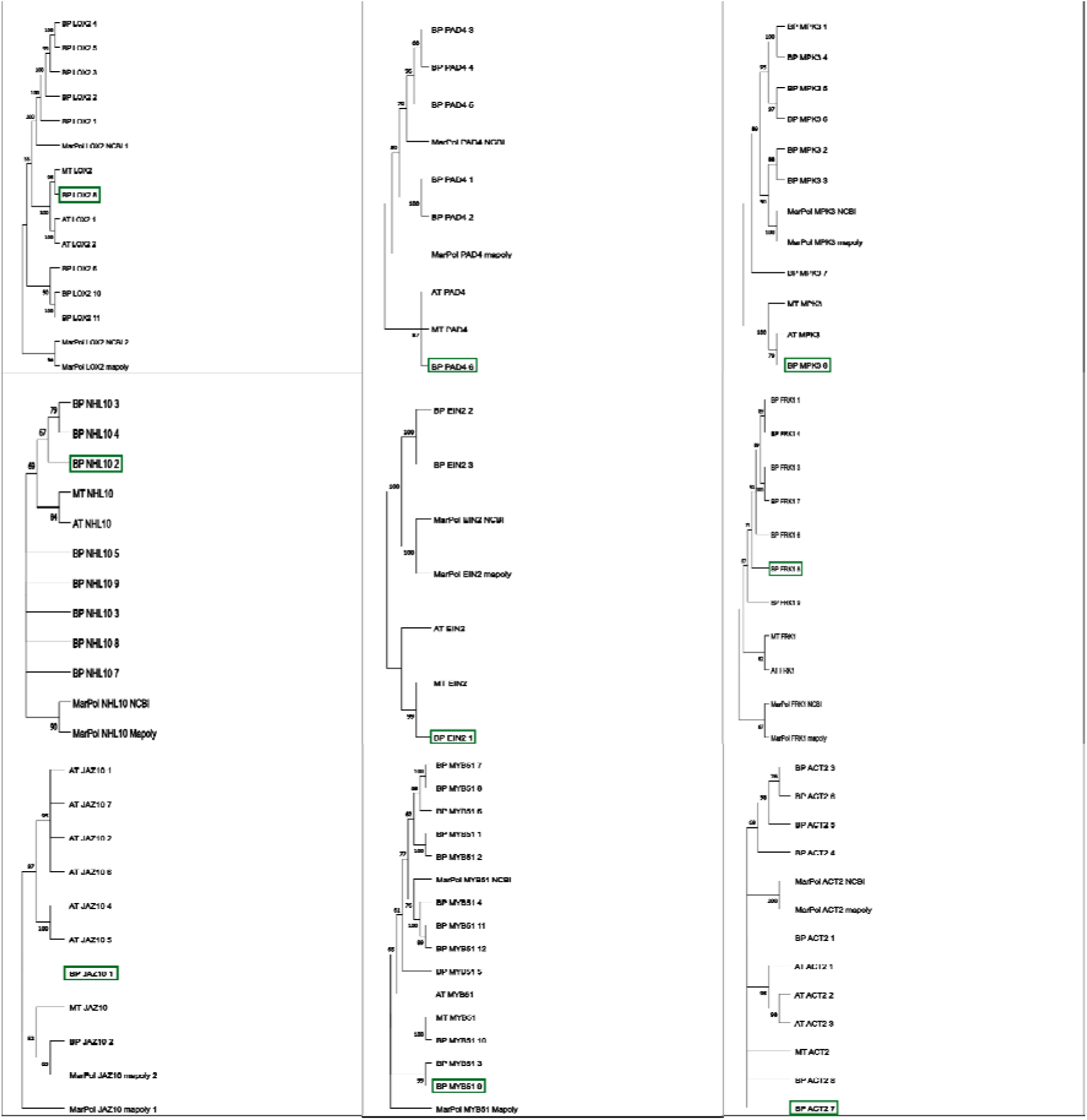
Cladograms of putative *Betula pendula* marker gene homologs. Phylogeny was based on translated protein sequences of *JAZ10, MYB51, EIN2, PAD4, LOX2, NHL10, FRK1, MPK3*, and reference gene *ACT2* from *Betula pendula* (BP), *Arabidopsis thaliana* (AT), *Medicago truncatula* (MT), *Marchantia polymorpha* (MarPol). In case of multiple spliced variants, all sequences were included for tree construction using the Maximum likelihood method. Green box denotes the homolog used for primer design in the study. In cases where multiple potential homologues clustered near MT and/or AT sequences, the candidate with the highest homology scores (provided by the BLAST tools used in each case) as well as with similar results in the reverse BLAST procedure were chosen.

**Supplementary Figure 3.**
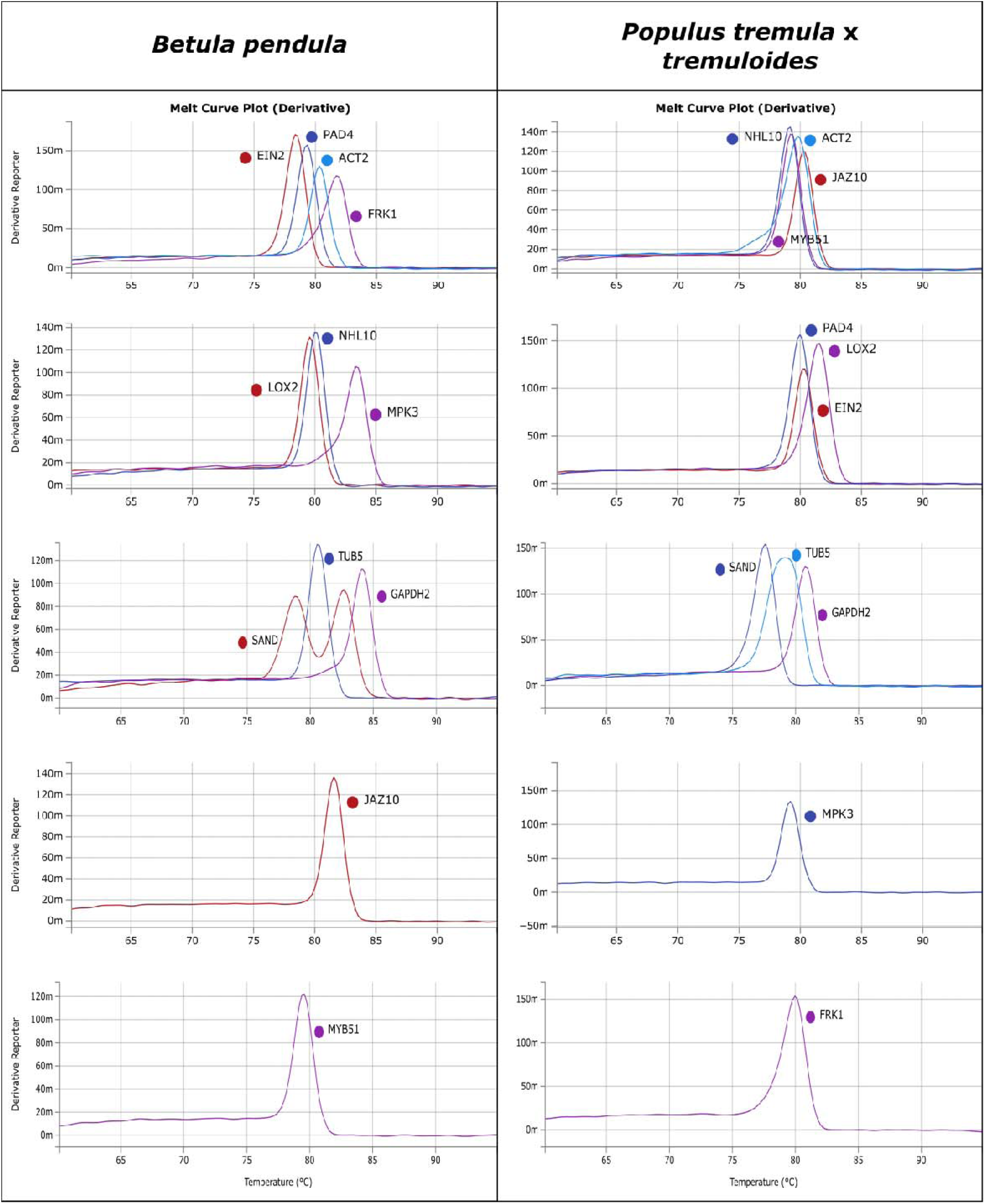
Melting curves for all tested gene amplicons in rt-qPCR data from representative samples of *Betula pendula* and *Populus tremula x tremuloides*.

**Supplementary Figure 4.**
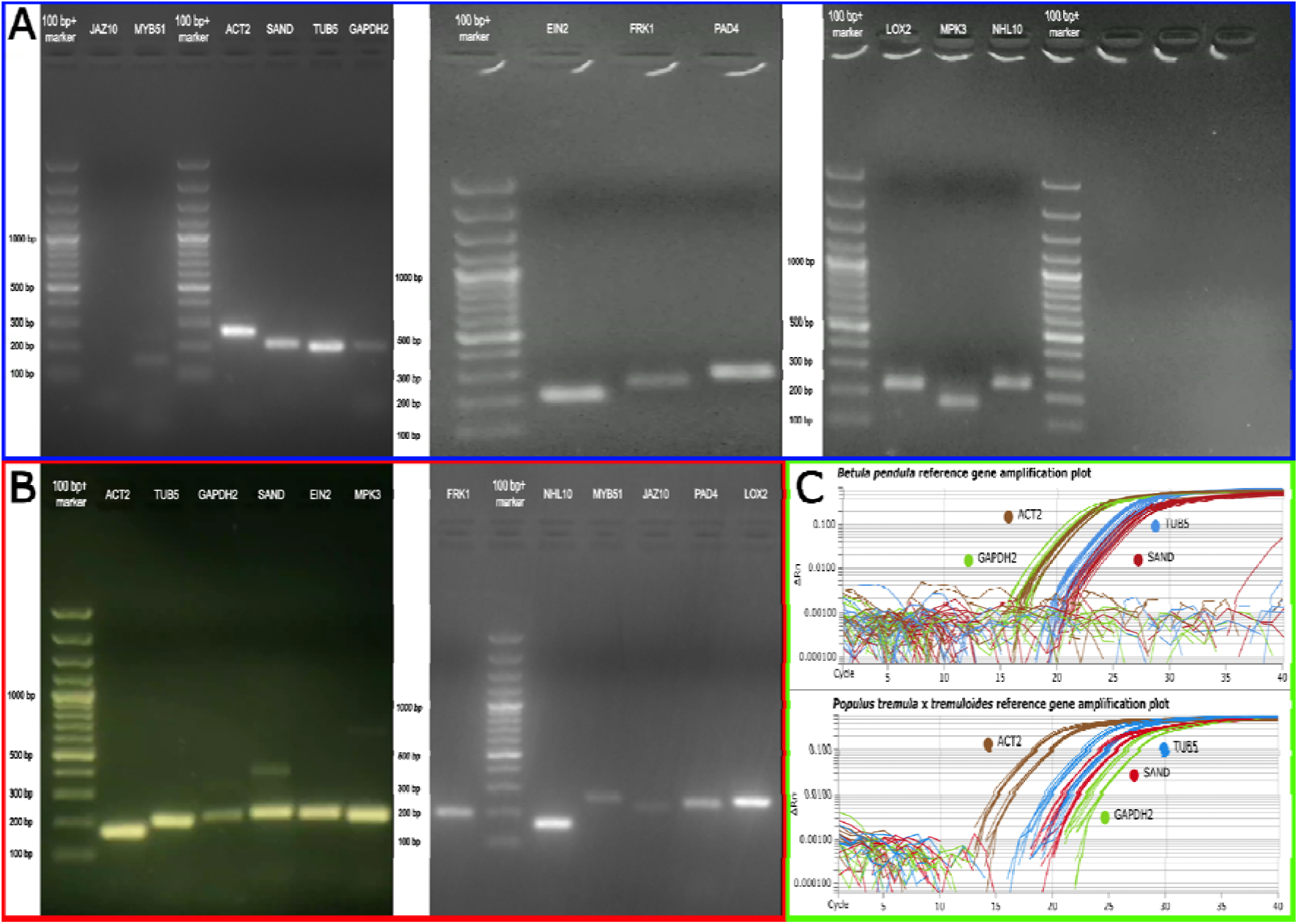
Electrophoresis gels of stress response gene amplicons in *Betula pendula* (A) and *Populus tremula x tremuloides* (B) as well as gene amplification curves (C) and variance across treatments (D) for tree reference gene selection. 35-cycle qPCR amplicons were used to check for primer specificity, target size and absence of double bands (**A-B**). deltaRn indicates the intensity of SYBR signal corresponding to target amplification during 35 qPCR cycles. Reference gene *ACT2* was selected based on earlier amplification and uniform expression across the different stress treatments in the qPCR amplification curve (**C**).

**Supplementary Figure 5.**
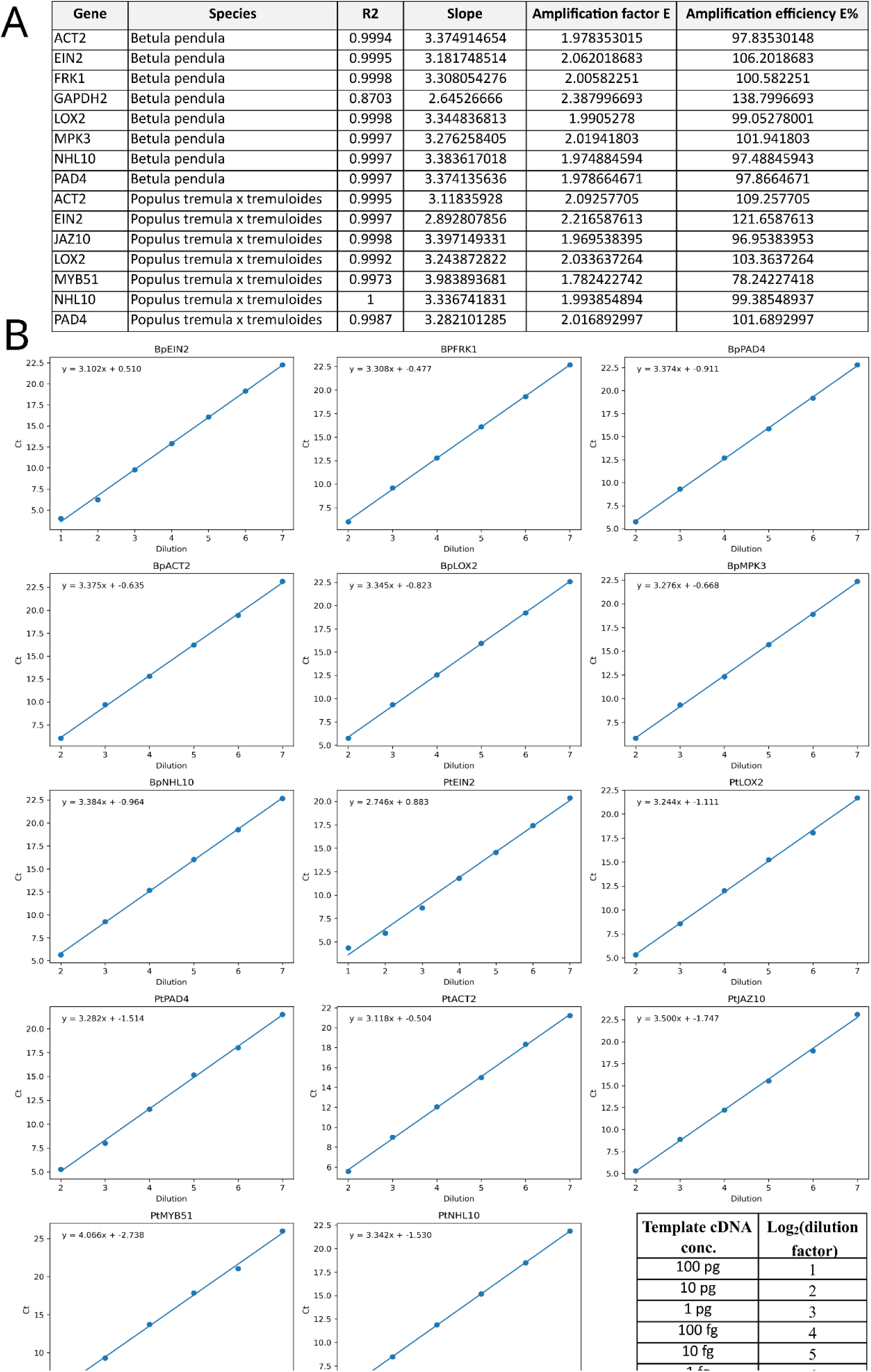
Calculation of primer efficiency for genes used in this study. **(A)** All stress marker genes used in this study are listed with their amplification efficiency (E%) and amplification factor (E) used for calculating ΔCt values. Calculation of E% is based on formula 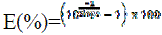. (**B**) Slope values are based on the regression lines for the Ct values for amplification of 10-fold dilution series of the template DNA, starting from 100 pg.

**Supplementary Figure 6.**
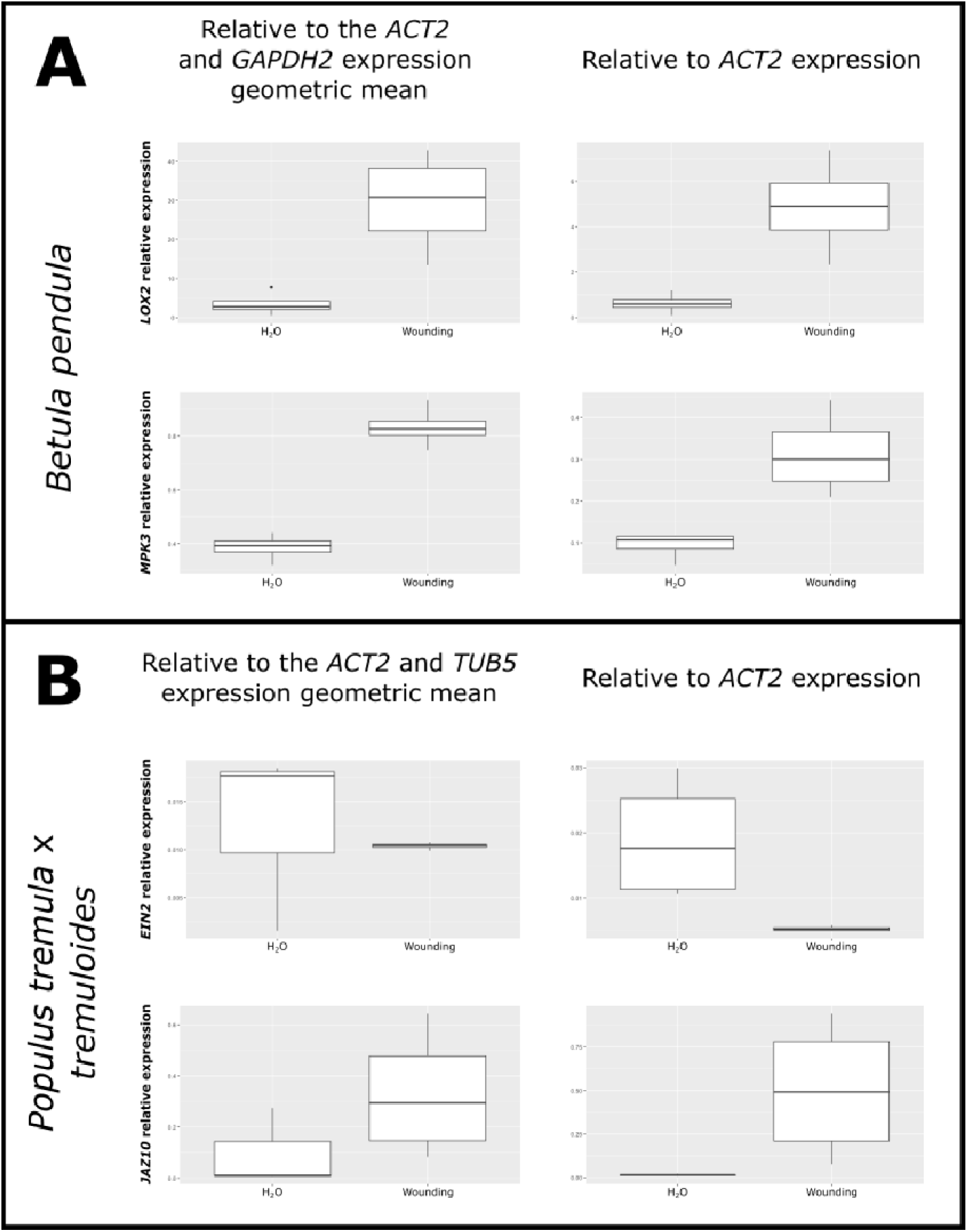
Reference gene testing for *B*. *pendula* (**A**) and *P*. *tremula* x *tremuloides* (**B**). To test whether *ACT2* is the optimal reference gene for defence gene expression testing, qPCR was carried out with an additional potential reference gene - *GAPDH2* and *TUB5* for *B*. *pendula* and *P*. *tremula* x *tremuloides* respectively and the relative expression of select genes and treatments was calculated relative to the geometric mean of both potential reference genes and compared to original expression data (only *ACT2* as a reference). Geometric mean of two selected reference genes is calculated with the formula 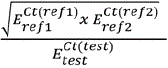, where *E_ref_*– primer efficiency of reference genes 1 and 2, *E_test_* – primer efficiency of test gene, *Ct(ref)* – Ct value of reference genes 1 and 2, *Ct(test)* – Ct value of the test gene. While both tested hybrid aspen genes showed differences in statistical significance, the expression tendencies remained similar in both cases. Difference in result significance could potentially be explained by the fact that the original models, which included all treatments, used ANOVA and/or Kruskal-Wallis tests, while the present experiment used t and Wilcoxon signed ranked tests, since only two groups were compared.

**Supplementary Figure 7.**
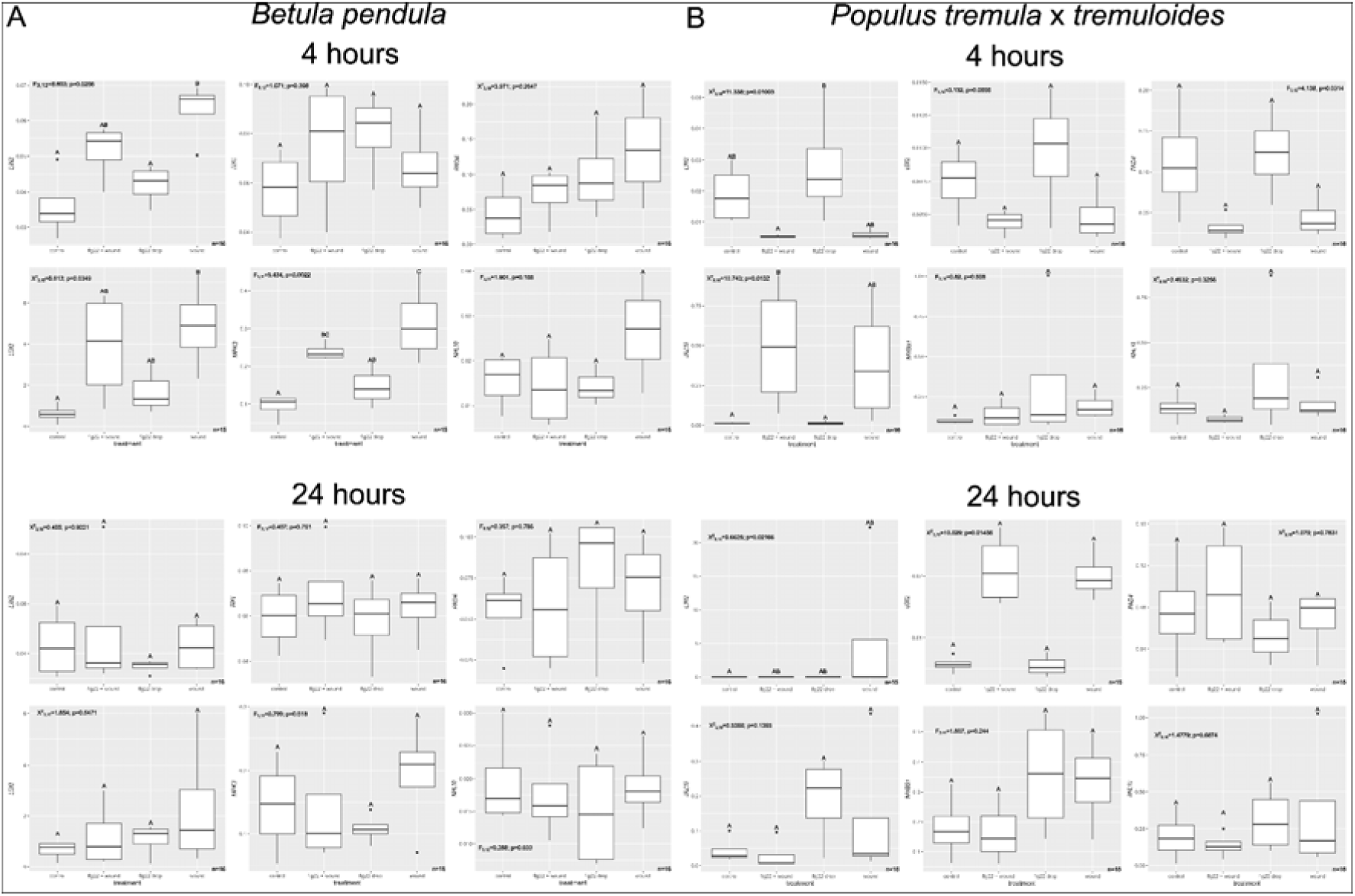
Marker gene expression relative to reference gene in *Betula pendula* (A) and *Populus tremula x tremuloides* (B). Boxplots represent the interquartile range (IQR), the line indicates the median expression, the whiskers - variance within 1.5x IQR. Summary results of ANOVA or Kruskal-Wallis and corresponding post-hoc tests are provided for each marker gene. Gene expression was normalized to ACT2 as reference. Ct were calculated as average from 3 technical replicate measurements for each sample. Test and reference gene was analyzed on the same qPCR plate. Each plate contained 4 independent biological replicates of at least 2 treatments. Sample size (n) for ANOVA or Kruskal-Wallis is shown at the bottom-left corner of each graph. Cases where sample size was 15 instead of 16, were related to insufficient cDNA amount.

**Supplementary Figure 8.**
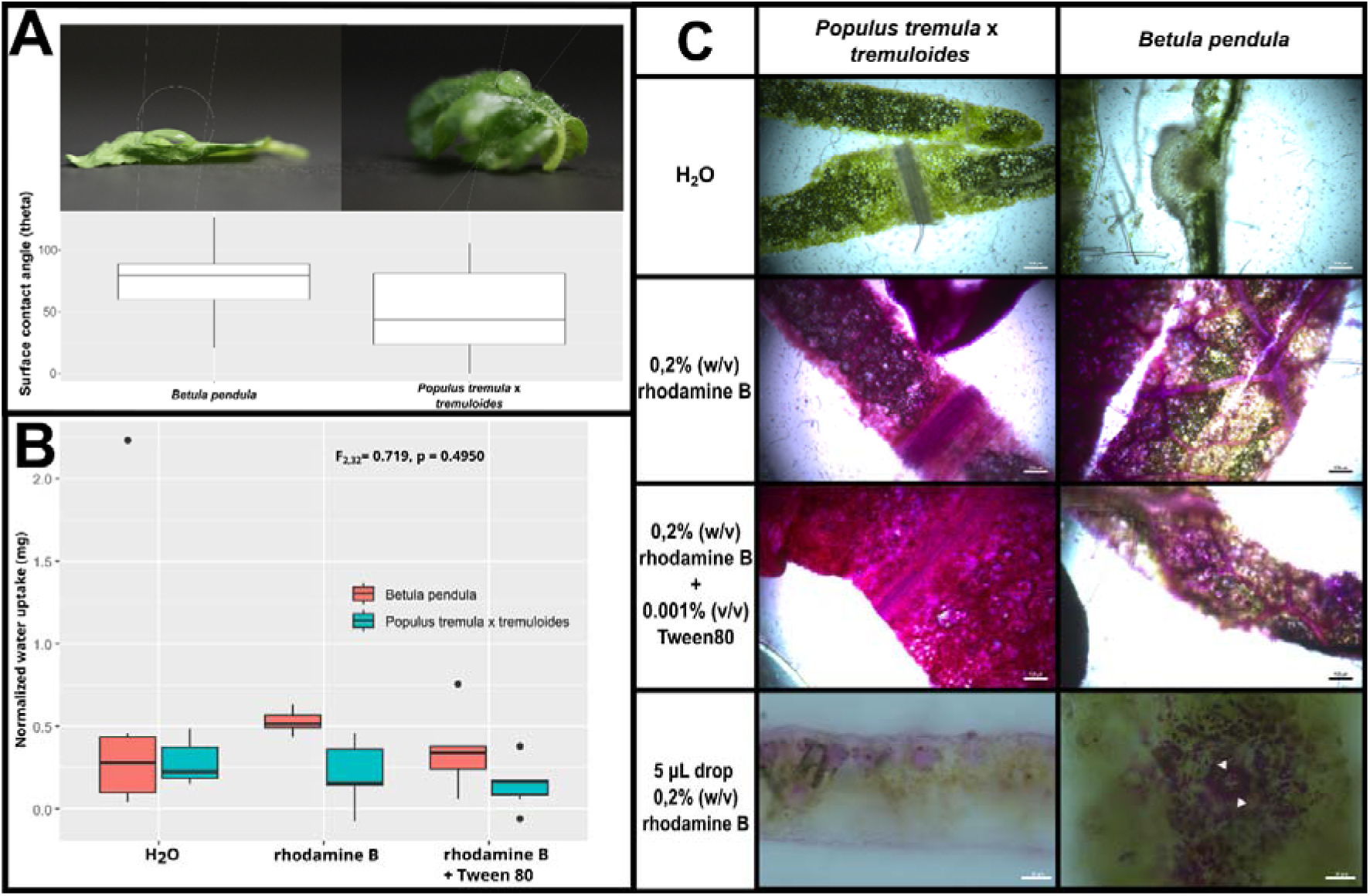
Silver birch and hybrid aspen leaves display leaf surface wettability, foliar water and dye uptake. (**A**) Leaves displaying droplet contact angle *theta* <90° are considered wettable according to Aryal & Neuner (2010). The wettability of the leaf surface for both tree species was determined by measuring the surface contact angle of a 5 μL 1μM flg22 drop on the surface of 12 independent leaves (representative images in the photographs). (**B**) No significant difference in water uptake 3 h after submergence was observed between both tree species. *B*. *pendula* displayed tendency for higher dye (0,2% w/v rhodamine B) uptake compared to *P*. *tremula* x *tremuloides*. Addition of 0.001% v/v surfactant Tween80 did not significantly improve dye solution uptake. The water or dye uptake was measured by weighing detached leaves as described in the methods section and based on Limm *et al*. (2009). (**C**) Light microscopy of *B*. *pendula* and *P*. *tremula* x *tremuloides* leaves 3h after submergence in water, rhodamine B solution or rhodamine B with added Tween 80 as well as application of a single 5 μL drop of rhodamine B solution on the abaxial leaf surface. Submergence photos (rows 1-3) were taken with 10x magnification (bar=100 μm). Rhodamine drop images (row 4) with 40x (bar=30 μm). White triangles on the *Betula pendula* photo indicates stomal guard cells - a potential entry point for rhodamine B solution into the leaf.

## Supplementary Files

**Supplementary File 1. Detailed description of the modified CTAB extraction protocol used for total RNA extraction.**

Prior to the extraction, all labware was treated with RNaseZap™ tissues to eliminate any traces of contaminants. The 1.5- and 2-mL test tubes were submerged in 0.1% DEPC solution for 24 h and autoclaved three times at 121 °C for 40 min, using saturated steam with pressure of at least 103 kPa (15 psi), to ensure the degradation of DEPC. Leaf samples were homogenized in liquid nitrogen using a pestle and mortar before transferring the material into 2 mL test tubes with 900 μl CTAB extraction buffer (2-cetyltrimethylammonium bromide; 0.1M Tris-HCl (pH 8); 1.4M NaCl; 20mM Ethylenediaminetetraacetic acid (EDTA) (pH 8); 2% polyvinylpolypyrrolidone (PVPP)) and 100 μl of β-mercaptoethanol). The mixture was shaken for 30 s and incubated in a thermoblock for 10 min at 65 °C. After incubation, 800 μl of chloroform were added to the mixture, which was followed by repeated shaking for 30 s. The mixture was then centrifuged for 10 min at 4 °C and 10’000 rpm.

The supernatant was transferred to a new 2 mL test tube together with 800 μl of a phenol/chloroform/isoamyl alcohol mixture (50:49:1) and shaken for 30 s. Samples were then centrifuged for 10 min at 10’000 rpm and 4 °C and the supernatant was transferred to a new 2 mL test tube together with an equal volume of a chloroform and isoamyl alcohol mix (24:1). The samples were then shaken for 30 s and centrifuged at 10’000 rpm at 4 °C. The supernatant was transferred to a new 1.5 mL test tube together with a third of the supernatant volume of 8M LiCl. Subsequently the samples were incubated for 24 h at -20°C, after which they were centrifuged for 20 min at 4 °C and 10000 rpm.

The sediment pellets were then washed with 1 mL 96% and 70% ethanol and centrifuged for 5 min at 10000 rpm. After centrifugation, ethanol was discarded, and the samples were dried at room temperature. The sediment pellets were then dissolved in 15 μl of 1x TE buffer with 1 μl of 1u/μl RNasin® Plus ribonuclease inhibitor solution (Promega). The mix was then treated with RQ1 Rnase free Dnase kit (Promega) according to manufacturer instructions. This process was followed by RNA extraction using a phenol/chloroform/isoamyl alcohol mixture (50:49:1). For convenience, 230 μl of 1x TE buffer were added to 10 μl of the sample. The phenol/chloroform/isoamyl alcohol mixture was added at a ratio of 1:1 to the sample solution.

After the extraction, RNA was sedimented by adding 0.1 volume of 3M sodium acetate and 2 volumes of 96% ethanol, which was followed by 24 hours of incubation at -20°C. Following the incubation period, the mixture was centrifuged for 15 min at 13’000 rpm and 4 °C and subsequently washed with 70% ethanol. The RNA sediment pellets were dried at room temperature and dissolved in 20 μl of TE buffer together with 1 μl of RNase inhibitor. The mRNA samples were stored at -80 °C. Total mRNA sample concentration and purity were measured using a “NanoDrop 2000” spectrophotometer (Thermo Scientific).

**Supplementary File 2. A representative selection of R studio code lines used in data analysis**

\# Libraries----

\# Data import

library(readxl) # For importing excel files

\# Statistics

library(car) # For Levene test

library(dunn.test) # For Dunn post-hoc test

\# Visualization

library(ggpubr) # For qqplots

library(qqplotr) # For qqplots

library(pheatmap) # For heatmaps

library(ggplot2) # For plot adjustments

\# Linear model and data manipulation----

library(plyr) # For data manipulation

library(emmeans) # For estimated marginal means

library(multcomp) # For comparisons

library(multcompView) # For compact letter displays

library(tidyverse) # For workflow

\# Data import----

data_bp <- read_excel(“qpcr_data_both_trees.xlsx”, sheet=1)

is.factor(data_bp$Treatment)

data_bp$Treatment <- as.factor(data_bp$Treatment)

data_bp_ein2_4h <- data_bp %>% # For group comparisons

filter(Gene==“EIN2”) %>%

filter(Timepoint==“4h”)

Heatmap_BP_dati_4h<- read_excel(“qpcr_data_both_trees.xlsx”, sheet=2) %>% # For heatmaps

filter(Timepoint==“4h”) %>%

group_by(Gene, Treatment) %>%

select(Treatment, Gene, Value) %>%

pivot_wider(names_from = Gene, values_from = Value)

\# Data vizualization----

bp_ein2_4h_boxplot <- ggplot(data_bp_ein2_4h,aes(Treatment, Value)) + geom_boxplot()

bp_ein2_4h_boxplot

\# Homogeneity of variance----

leveneTest(Value ∼ Treatment, data = data_bp_ein2_4h)

\# Group comparison - ANOVA----

ein2_4h_aov <- aov(Value ∼ Treatment, data = data_bp_ein2_4h)

summary(ein2_4h_aov)

\# Checking the distribution of residuals----

ggplot(data.frame(resid = residuals(ein2_4h_aov)), aes(sample = resid)) +

geom_qq_band() + stat_qq_line() + stat_qq_point() +

labs(x = “Theoretical quantiles”, y = “Sample quantiles”)

shapiro.test(residuals(ein2_4h_aov))

\# Post hoc test----

ein2_tukey <- TukeyHSD(ein2_4h_aov)

ein2_tukey

\# If residuals were non-normal - Kruskal-Wallis test----

ein2_4h_krusk <- kruskal.test(Value∼Treatment, data = data_bp_ein2_4h)

ein2_4h_krusk

\# Alternative post-hoc test - Dunn test---

ein2_dunn <- dunn.test(data_bp_ein2_4h$Value, data_bp_ein2_4h$Treatment, method = “bonferroni”)

\# Heatmap----

BP4=as.matrix(Heatmap_BP_dati_4h[,-1])

row.names(BP4)=Heatmap_BP_dati_4h$Treatment

BP4_heatmap <- pheatmap(BP4, fontsize=14, main = “BP 4 hours”)

BP4_heatmap

\# Linear model----

birch<-as.data.frame(read_xlsx(“qpcr_data_both_trees.xlsx”, sheet=1)) #additional column specifying species should be added

aspen<-as.data.frame(read_xlsx(“qpcr_data_both_trees.xlsx”, sheet=3)) #additional column specifying species should be added

data<-rbind(birch, aspen)

rm(birch, aspen)

data$id<-paste(data$Species,data$Gene, data$Timepoint, sep=“_”)

contr<-ddply(data, .(id), summarize,

control_mean=mean(Value[Treatment==“Control (H2O drop)”]))

data<-join(data,contr, by=“id”, type=“left”)

rm(contr)

data$logfold1<-log(data$Value/data$control_mean,2)

logfold<- data[data$Treatment1!=“Control (water)”,]

mod<-lm(logfold∼Treatment*Gene*Timepoint,

offset = control_mean,

data = logfold[logfold$Species==“Silver birch”,])

## Notes

### Competing Interest Statement

The authors have declared no competing interest.

### Summary of Updates

We have made numerous and substantial improvements in the main text, figures and their legends as well as supplemental material. For example, we have performed additional experimental work to determine leaf wettability and in silico analysis on promoter divergence of stress marker gene LOX2, as suggested by the reviewer. This resulted in additional main and supplemental figures. Moreover, we have streamlined text, improved on clarity of key terms, and the coherence of the manuscript's structure overall.

### Funding

- Latvian Council of Science Fundamental and Applied Research: lzp-2022/1-0283

