## Supplementary figures and images for "Species specific marker genes for systemic defence and stress responses to leaf wounding and flagellin stimuli in hybrid aspen and silver birch"

### Supplemental Figure 1

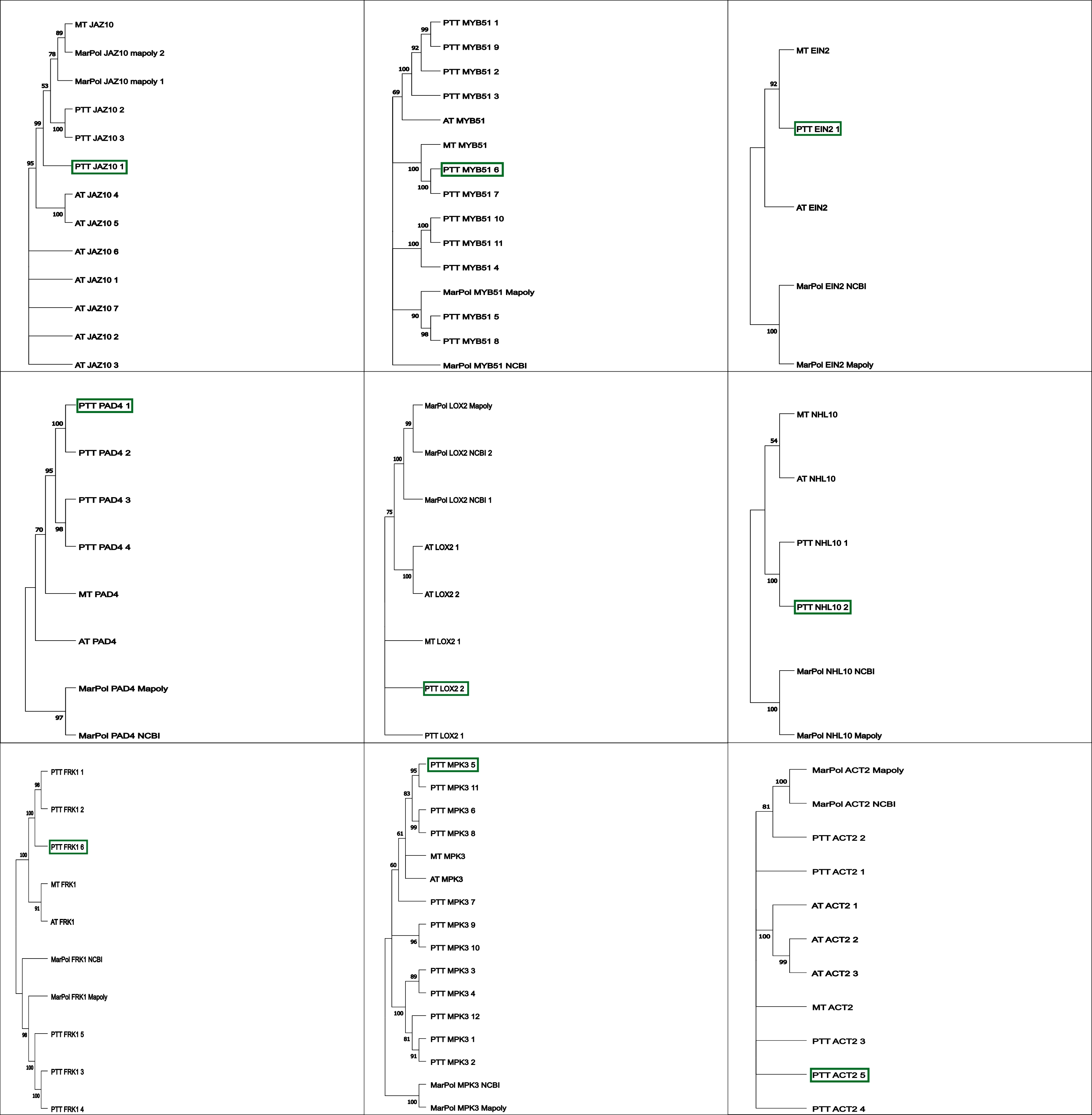

### Supplemental Figure 2

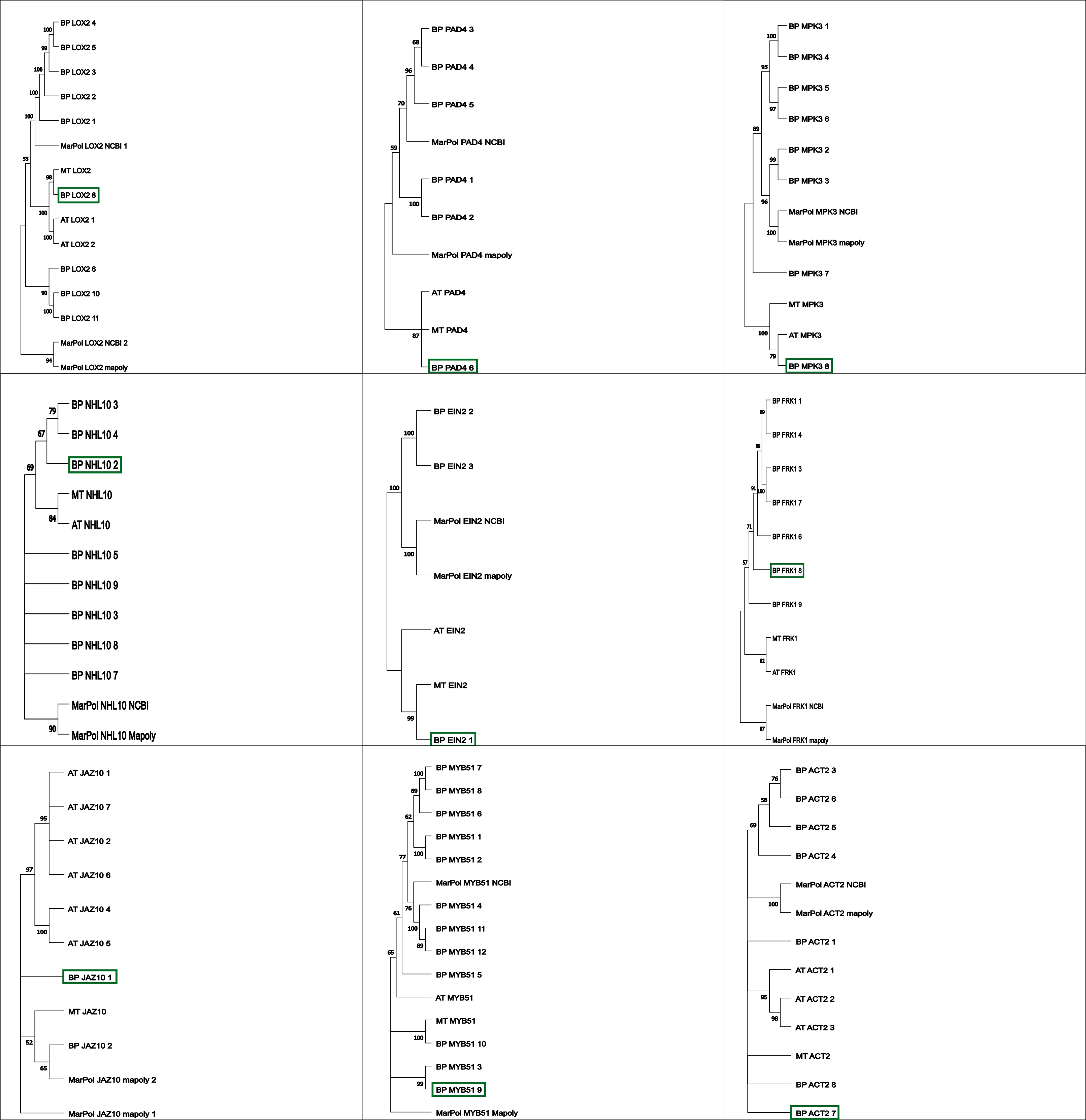

### Supplemental Figure 3

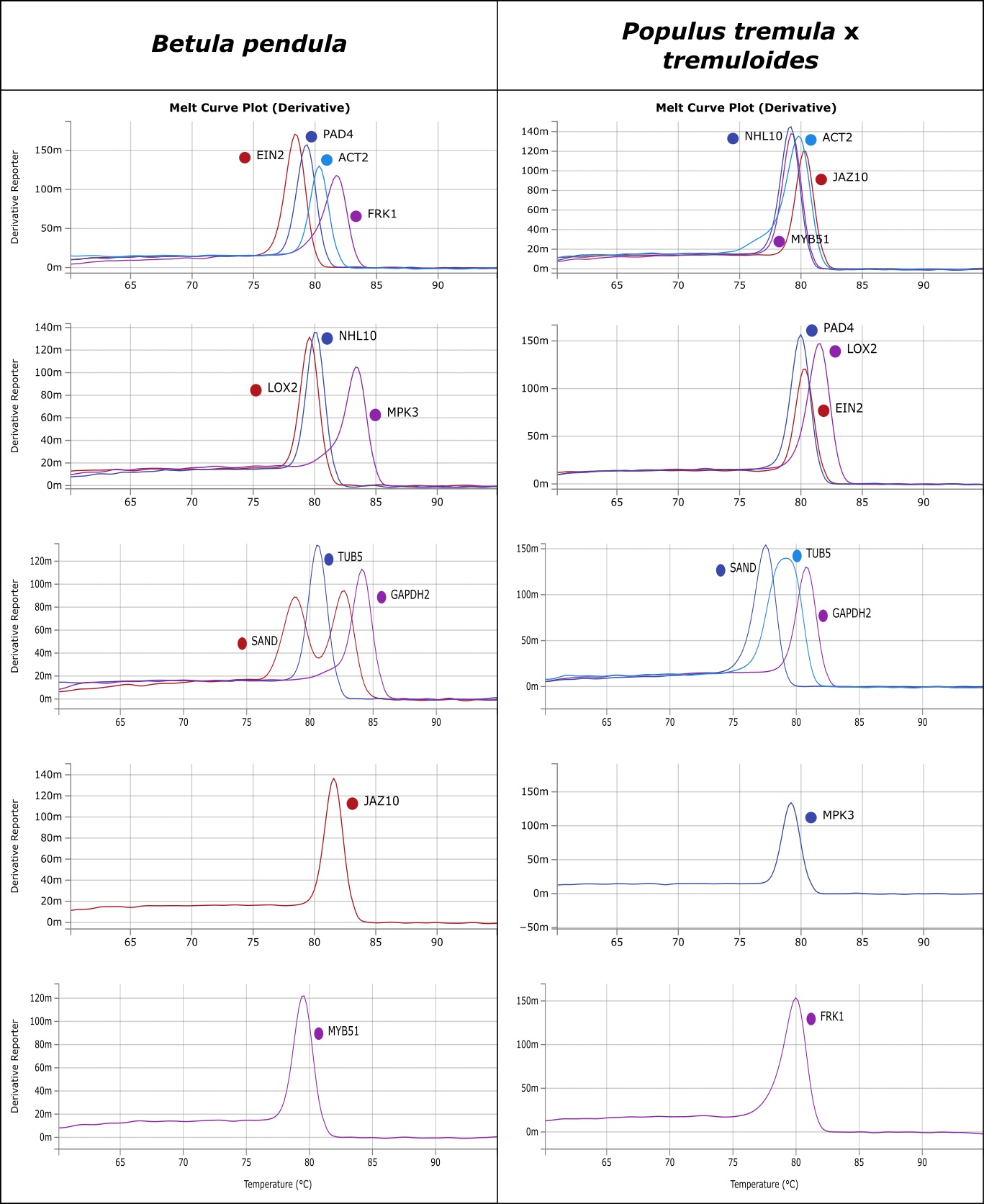

### Supplemental Figure 4

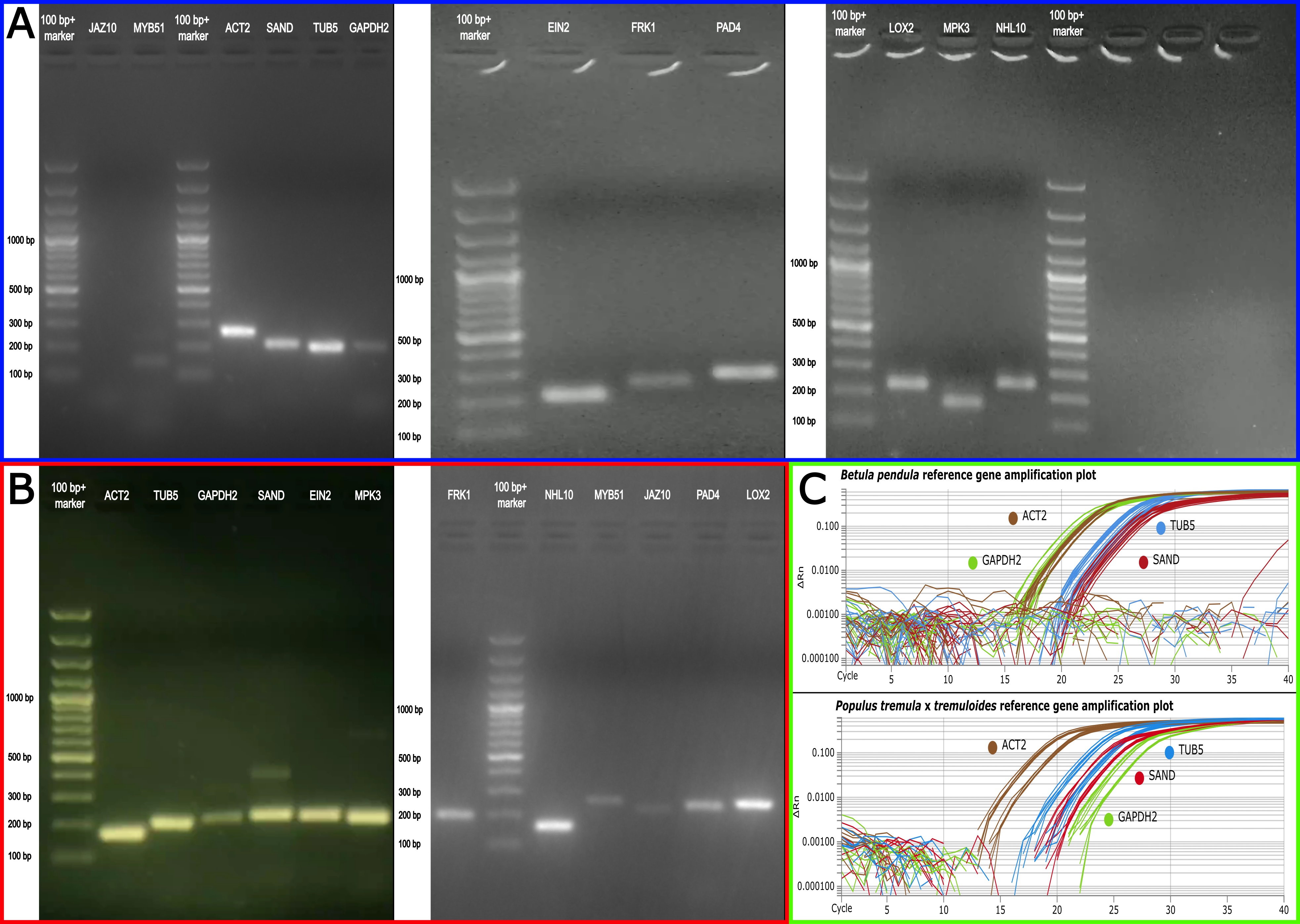

### Supplemental Figure 5

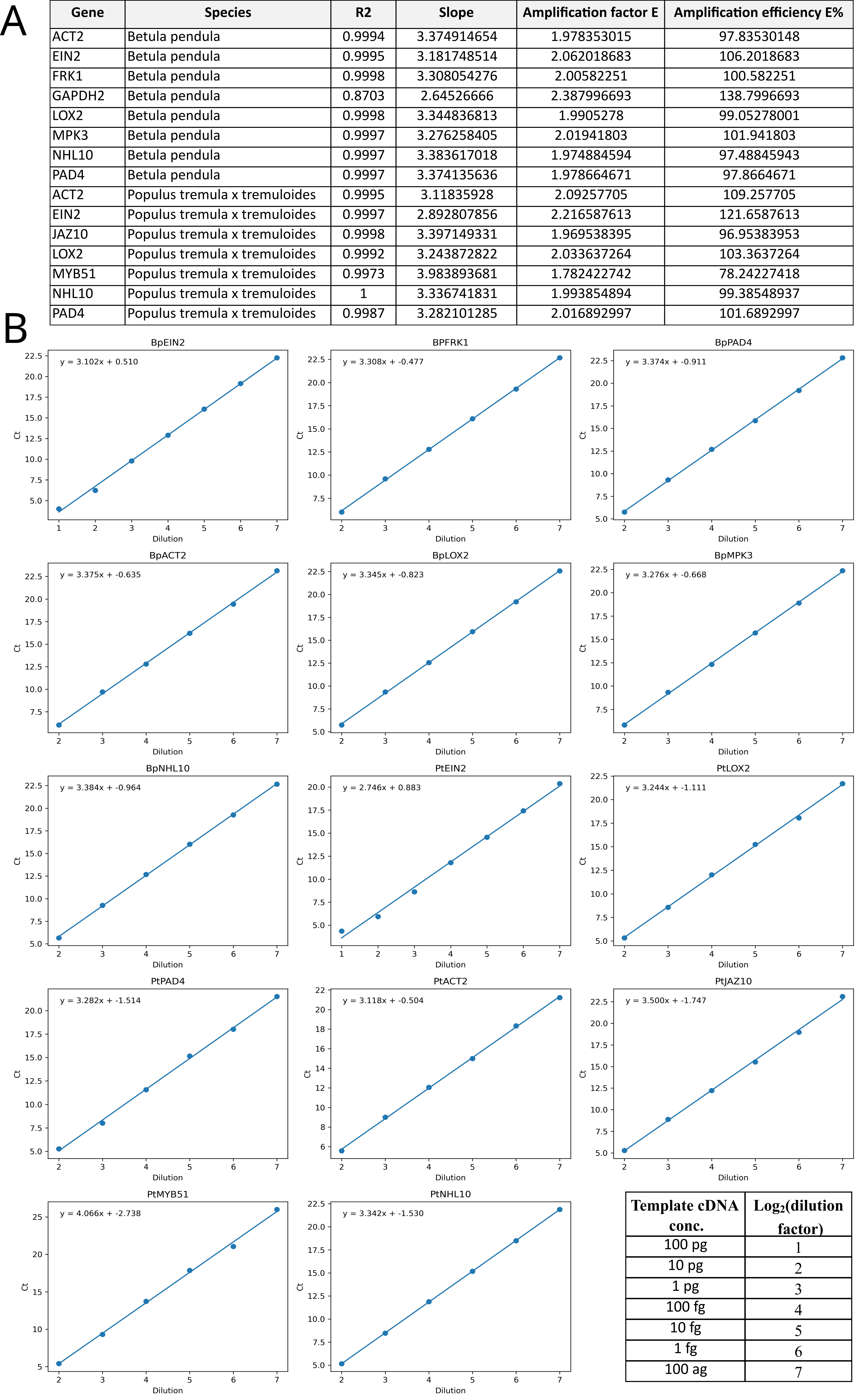

### Supplemental Figure 6

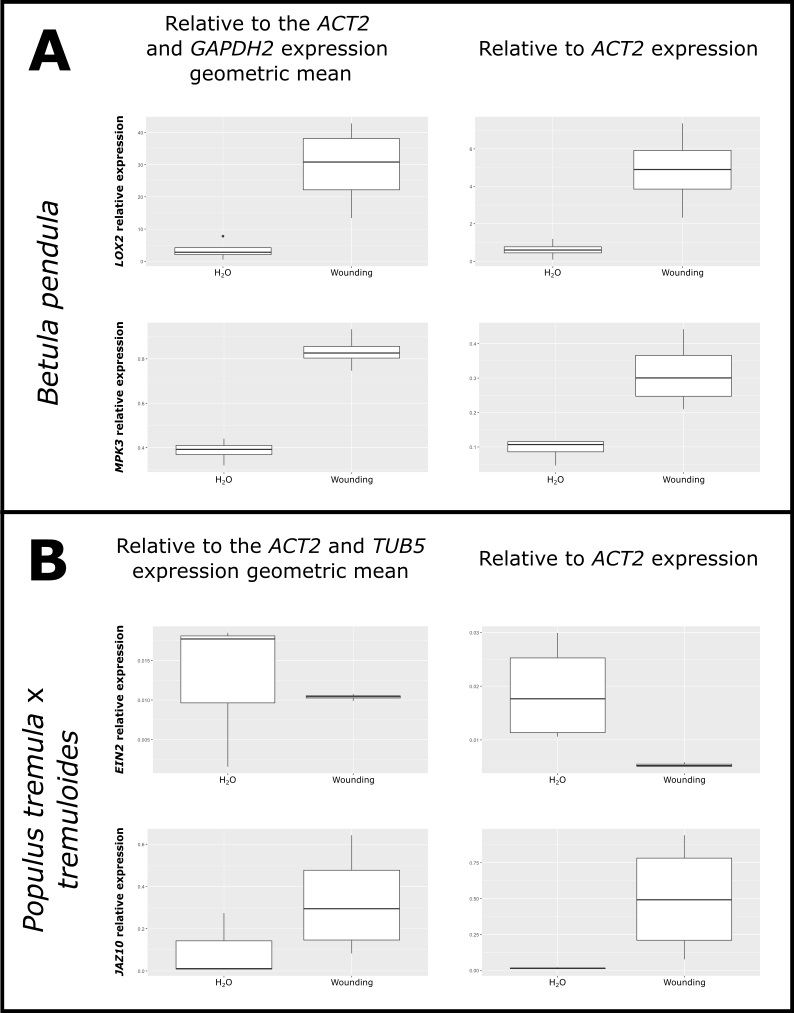

### Supplemental Figure 7

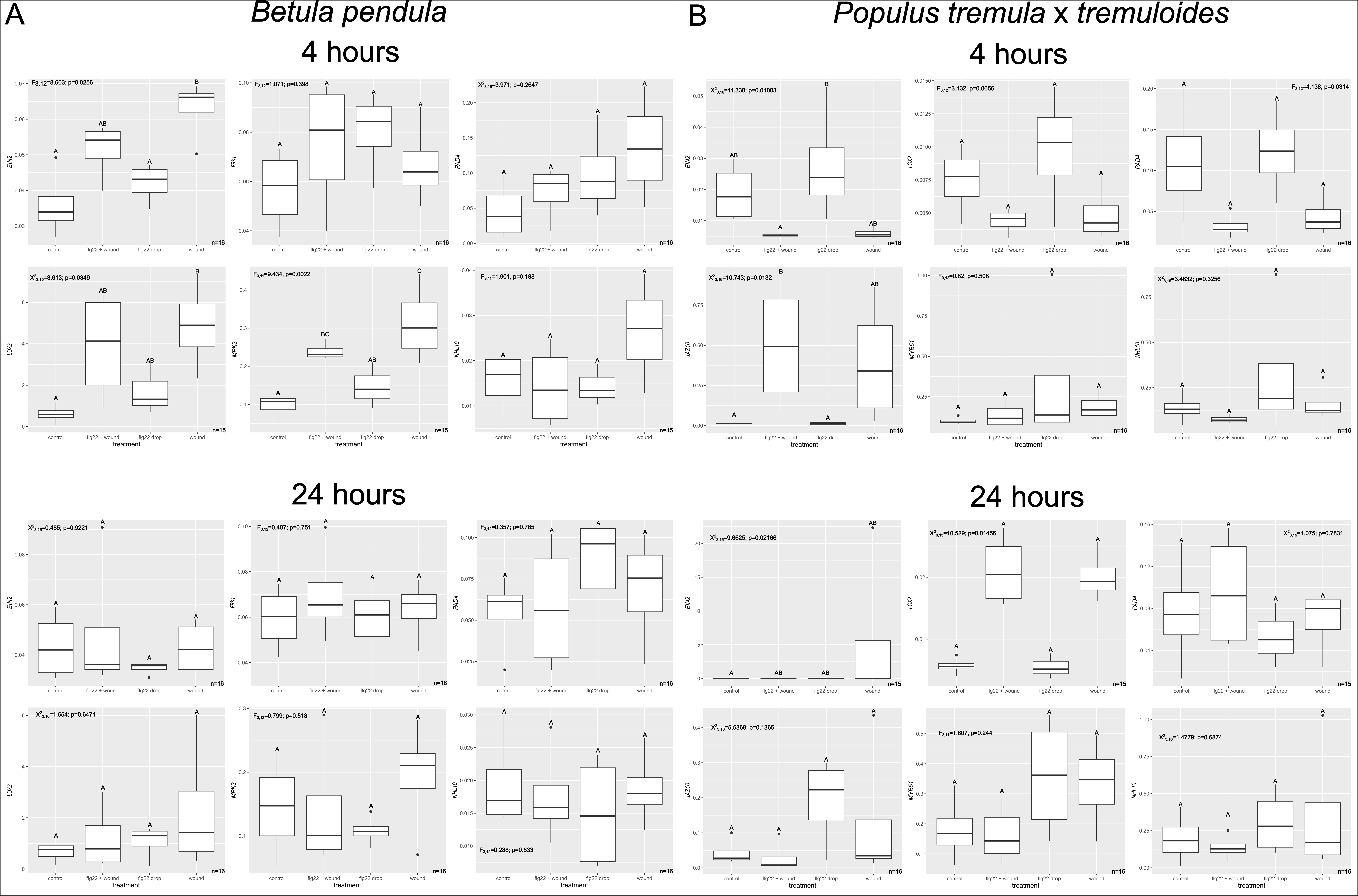

### Supplemental Figure 8

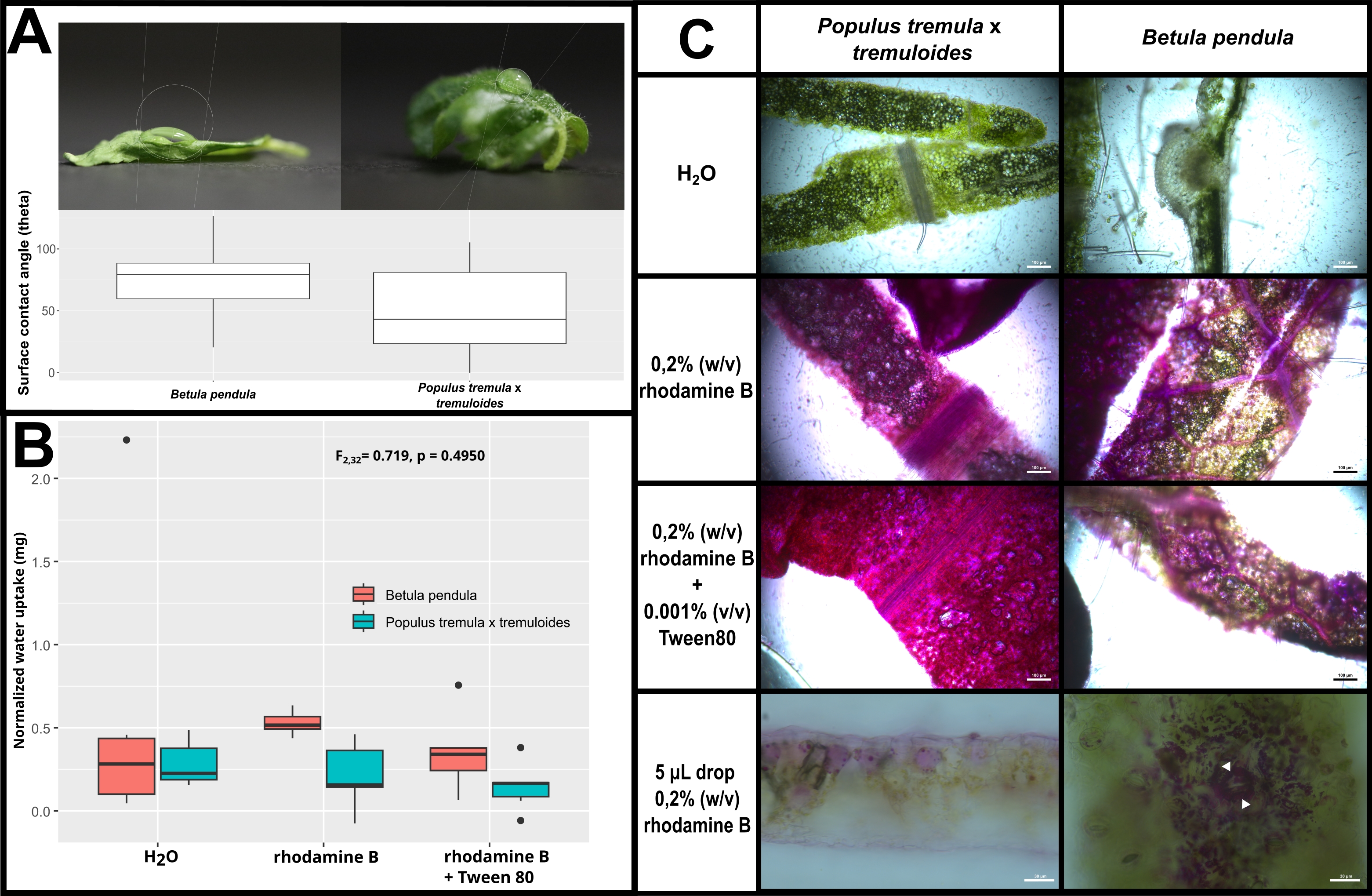
